# Structures of RecBCD in complex with phage-encoded inhibitor proteins reveal distinctive strategies for evasion of a bacterial immunity hub

**DOI:** 10.1101/2022.09.12.506733

**Authors:** M. Wilkinson, O.J. Wilkinson, C. Feyerherm, E.E. Fletcher, D.B. Wigley, M.S. Dillingham

## Abstract

Following infection of bacterial cells, bacteriophage modulate double-stranded DNA break repair pathways to protect themselves from host immunity systems and prioritise their own recombinases. Here we present biochemical and structural analysis of two phage proteins, gp5.9 and Abc2, which target the DNA break resection complex RecBCD. These exemplify two contrasting mechanisms for control of DNA break repair in which the RecBCD complex is either inhibited or co-opted for the benefit of the invading phage. Gp5.9 completely inhibits RecBCD by preventing it from binding to DNA. The RecBCD-gp5.9 structure shows that gp5.9 acts by substrate mimicry, binding predominantly to the RecB arm domain and competing sterically for the DNA binding site. Gp5.9 adopts a parallel coiled-coil architecture that is unprecedented for a natural DNA mimic protein. In contrast, binding of Abc2 does not substantially affect the biochemical activities of isolated RecBCD. The RecBCD-Abc2 structure shows that Abc2 binds to the Chi-recognition domains of the RecC subunit in a position that might enable it to mediate the loading of phage recombinases onto its single-stranded DNA products.

## Introduction

The RecBCD complex of *E. coli* plays a dual role in bacterial cell biology, acting both as a phage immunity hub and as the primary mechanism to initiate the recombinational repair of double-stranded DNA breaks (DSBs) (1–3). This is facilitated by RecBCD’s ability to switch between different modes of action at free DNA ends. In the destructive mode, RecBCD recognises free DNA ends and acts as a processive exonuclease that degrades both strands of the DNA duplex. This is detrimental to bacteriophage for multiple reasons (1, 2, 4); it can directly degrade and inactivate linear phage genomes upon entry into the bacterial cell, it provides acquired immunity against future phage infections because degraded phage sequences are catalogued in CRISPR libraries, and it prevents phage replication in the rolling circle mode. However, RecBCD can also act in a recombinogenic mode to convert DSBs into 3’-terminated ssDNA overhangs, onto which it loads RecA protein to promote strand exchange. This facilitates DNA break repair which also underpins replication and protects the genetic stability of the host genome. The switch from the destructive to the recombinogenic mode is triggered during RecBCD translocation along DNA by recognition of a specific octameric ssDNA sequence called Chi (crossover hotspot instigator; 5’-GCTGGTGG-3’) (5). Chi sequences are highly over-represented in host genomes but are often lacking in bacteriophage DNA. Therefore, regulation of RecBCD by Chi has been viewed as a form of self-recognition which instructs RecBCD to repair host DNA but to destroy and catalogue foreign DNA.

Bacteriophage could evade the immunity function of RecBCD in several different ways (4). One strategy is to select for Chi sequences within their genome, but this is thought to incur a significant fitness penalty: phage are themselves dependent on recombination for packaging and genetic variation, and the use of the RecBCD pathway may provide sub-optimal outcomes and limit the breadth of possible hosts. Instead, many phage express inhibitors of RecBCD which enable them to avoid genome degradation and prioritise the use of their own autonomous recombination system; the phage-encoded exonuclease (*exo*) and recombinase (*bet*) genes. A well-characterised example of this strategy is provided by coliphage λ Gam protein. Gam acts as a DNA mimic protein to bind directly to RecBCD and inhibit all of its activities by competing for the DNA binding site (6–10). Analogous RecBCD inhibitors are also present in phage T4, T7 and SPP1 but these do not share sequence homology with Gam and are less well-characterised in comparison (11). The T7 gp5.9 protein, which is one subject of this work, was discovered as a protein responsible for reduced ATP-dependent exonuclease activity in extracts from cells infected with T7, and was subsequently shown to reduce the activity of purified RecBCD *in vitro* (11, 12). Despite the lack of primary structure homology to λ Gam, gp5.9 is also a small and highly acidic protein, both of which are properties associated with DNA mimicry (13, 14). Therefore, T7 gp5.9 may act via a similar mechanism to Gam by competing for the RecBCD DNA binding locus. Remarkably, bacterial strains containing the Ec48 retron system have turned this RecBCD evasion strategy to their own advantage (3). Within such cells, RecBCD interaction with Gam or gp5.9 acts as a sensor of phage infection and triggers cell suicide, protecting the whole bacterial population at the expense of infected cells.

A completely different strategy to evade the anti-phage activities of RecBCD is exemplified by the Abc2 protein of *Salmonella* phage P22 (9, 15–18). Abc2 binds to the RecC subunit, from where it is proposed to hijack RecBCD to promote recombination (rather than degradation) of phage. At the molecular level, this might be achieved by promoting the conversion of RecBCD into its recombinogenic mode and adapting the RecA-loading function for the phage-recombinase. This possibility is supported by several observations: phage encoding Abc2-like proteins apparently lack an exonuclease function (which would instead be provided by the co-opted RecBCD complex), Abc2 expression eliminates the SOS response to DSBs which requires RecA loading, and Abc2-RecBCD fails to respond to Chi yet displays properties somewhat similar to the Chi-modified enzyme constitutively (i.e. upregulated 5’-nuclease activity).

In this work, we present a comparative study of the interactions between RecBCD and the phage-encoded gp5.9 and Abc2 proteins. We find that there are stark differences in the effects of the two proteins on RecBCD activity. Whereas gp5.9 inhibits all tested activities of RecBCD, Abc2 has no significant effect on the ability of the complex to bind and unwind DNA. Accordingly, cryo-EM structures of gp5.9-RecBCD and Abc2-RecBCD show that the two phage proteins bind to entirely different parts of the RecBCD structure. The gp5.9 protein uses DNA mimicry to bind the RecB arm domain and compete for the interaction between RecBCD and DNA ends. The Abc2 protein, together with a host encoded prolyl-isomerase, binds to the RecC subunit from where it might influence the recombinase loading step of homologous recombination.

## Materials and Methods

### Protein expression and purification

The T7 gp5.9 gene from phage T7 was produced as a synthetic gene construct with an N-terminal 3C-cleavable histidine tag (GeneArt, Invitrogen). This was subcloned into the pACEBAC1 (MultiBac) vector using engineered BamHI and XbaI sites for overexpression in insect cells using standard techniques (19, 20). Briefly, for large scale expression of gp5.9, 500 mL of High5 cells at 2×10^6^ cells/mL were infected with 25 mL P3 virus and incubated for 72 hours at 27°C with shaking before cells were harvested by centrifugation. The pellet was resuspended in 50 mL lysis buffer (20 mM Tris pH7.5, 200 mM NaCl, 2 mM β-mercaptoethanol, 10% glycerol, protease inhibitor cocktail (Roche, as directed by manufacturer), 20 mM imidazole). The cells were lysed by sonication and centrifuged to remove cell debris. The supernatant was then applied to Talon resin (Takara Bio) to purify gp5.9 using the histidine tag. Beads were equilibrated by washing three times with 15 mL wash buffer (20 mM Tris pH 7.5, 200 mM NaCl, 20 mM β-mercaptoethanol, 10% glycerol, 20 mM imidazole). Supernatant from the centrifuged cell lysate was added to the beads and incubated for 30 minutes at 4°C. The beads were then spun down and the supernatant (unbound protein) was removed. The beads were washed 4 times with wash buffer before gp5.9 was eluted with 50 mL elution buffer (20 mM Tris pH 7.5, 200 mM NaCl, 20 mM β-mercaptoethanol, 150 mM imidazole). The eluate was loaded onto a 1 mL MonoQ column (GE Healthcare) in buffer A (20 mM Tris pH 7.5, 1 mM TCEP) and eluted with buffer B (20 mM Tris pH 7.5, 1 mM TCEP, 1 M NaCl). Peak fractions were pooled and further purified using size exclusion chromatography in SEC buffer (20 mM Tris pH 8, 0.5 mM TCEP, 200 mM NaCl). The concentration of his-tagged gp5.9 was calculated using a theoretical extinction coefficient of 8480. The final protein preparation was stored at −80°C in a buffer containing 20 mM tris-Cl pH 8, 0.5 mM TCEP, 200 mM NaCl at a final concentration of 19.4 μM. A sample of tag-free gp5.9 was prepared by treatment of this stock with 3C protease (Pierce) followed by size exclusion chromatography and the resulting protein was stored in the same buffer at a final concentration of 14.9 μM. Unless stated otherwise, all experiments presented in this paper were performed with the tag-free gp5.9 protein.

For the wild type RecBCD and nuclease-dead RecB^D^CD complexes, RecB, RecC and RecD were co-expressed and purified from three separate plasmids: pETduet-His_6_-TEVsite-recB or pETduet-His_6_-TEVsite-recB^D1080A^, pRSFduet-recC and pCDFduet-recD as described previously (21, 22). For the RecBCD-Abc2 complex, the four genes were co-expressed together as Abc2 has been shown to be unstable when overexpressed alone (15). The nuclease-dead RecB mutant (D1080A) was used to prevent digestion of the DNA substrates. To avoid adding a fourth plasmid to the transformation, the genes encoding His_6_-TEVsite-recB^D1080A^ and RecC were jointly cloned into the respective NdeI/KpnI and NcoI/XhoI sites of the pRSFduet vector for use with the pCDFduet-recD plasmid. Finally, the Abc2 gene from phage P22 was synthesised (GeneStrings, Thermofisher) and cloned in-between the NdeI and XhoI sites of pET22b. An Abc2^P68A^ mutant variant was generated using In-Fusion cloning (Takara-Bio) for the generation of RecBCD-Abc2 complex without the co-purification of the host PpiB protein. Co-expression of the three plasmid system coding for RecB, RecC, RecD and Abc2 was performed as for wild type RecBCD. The RecBCD-Abc2 complexes were purified by a similar method to that described previously for RecBCD (22). Briefly, the cells were lysed using an emulsiflex cell disruptor, the lysate clarified by centrifugation and ammonium sulphate was added to the soluble fraction (0.35 g/mL) before a second centrifugation at 30,000g to pellet the proteins. The pellets were resuspended and complexes bound to a HisTrap column (GE healthcare) before direct elution onto a HiTrap heparin column (GE healthcare) and subsequent elution using a gradient of NaCl. The His-tags were cleaved by incubation with TEV protease during overnight dialysis before being re-passed through the HisTrap column and finally eluted from a MonoQ column with a shallow gradient of NaCl. The purified proteins were diluted to give a final buffer solution of 25 mM Tris pH 7.5, 100 mM NaCl, 0.5 mM TCEP. The samples were concentrated to around 5-10 mg/mL, supplemented with glycerol to a concentration of 15 % (v/v) and flash-frozen in aliquots in liquid nitrogen for storage at −80°C. The fluorescent SSB biosensor was a gift from Martin Webb (The Crick Institute, London) and was purified as described (23).

### Preparation of DNA substrates for cryoEM and native gel shift assays

A splayed hairpin DNA substrate as used in previous cryo-EM studies (22) was also used here to allow direct comparison of the effect of Abc2 interaction on the conformation of the RecBCD-DNA complex. The oligonucleotide sequence is 5’-TTT TTT TTT TTT tct aat gcg agc act gct aca gca tTT CCC atg ctg tag cag tgc tcg cat tag aTT T −3’, with lower case denoting paired bases in the duplex region and upper case denoting unpaired bases (twelve on the 5’-end and three on the 3’-end). The DNA substrate was purified as described previously (21, 24). Briefly, the synthesised oligonucleotide (IDT) was annealed at low concentration by heating to 95°C followed by a slow cooling to room temperature and purification by chromatography on a Source-Q column. The purified substrate was desalted into deionised water, aliquoted and stored at −20°C.

### DNA double-strand break resection assays

DSB resection assays were performed in RecBCD buffer (25 mM Tris pH7.5, 10 mM NaCl, 6 mM MgCl_2_, 0.1 mg.ml^−1^ BSA) supplemented with 2.5 nM RecBCD (or RecBCD-ABC2 or RecBCD-ABC2-PPI) or 20nM AddAB, 960 μM (in ntds) linearised pACEBac1 vector, and 1 μM gp5.9. Reactions were initiated by the addition of 2 mM ATP. 10μL aliquots were removed at the times indicated and quenched by the addition of 10μL STOP buffer (10% glycerol, 1% SDS, 50mM EDTA, 1mg/mL proteinase K). The samples were run on 1% agarose gels and imaged by post-staining with SYBR gold.

### Helicase assays

Real time helicase assays were performed using a fluorescent biosensor for ssDNA based on the *Plasmodium falciparum* SSB protein (fSSB; (23)). Reactions were performed in RecBCD buffer (25 mM Tris pH7.5, 10 mM NaCl, 6 mM MgCl_2_, 0.1 mg.ml^−1^ BSA) supplemented with 10 pM RecBCD, 1 uM (in ntds; ~100 pM molecules) bacteriophage lambda DNA (New England Biolabs), 25 nM fSSB (tetramer) and gp5.9 at the stated concentration. After a 10 minute pre-incubation, reactions were initiated by the addition of 2 mM ATP. Fluorescence intensity was monitored using a Cary Eclipse Fluorescence Spectrophotometer (excitation wavelength 430 nm, emission wavelength 475 nm, excitation and emission slit widths of 10 nm and 5 nm respectively). Assays were performed in triplicate and the initial rates reported are the mean and standard error for the three repeats. To obtain IC_50_ values for inhibition of RecBCD by gp5.9, data describing the initial unwinding rate as a function of log_10_[gp5.9] were fit to the sigmoidal dose response equation of GraphPad Prism.

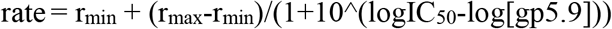

r_min_ and r_max_ are the minimum and maximum DNA unwinding rates respectively. Values for the unwinding rates were normalised to 100% for a zero gp5.9 control and so for the purposes of the fits the values for r_max_ and r_min_ were constrained to 100 and 0 respectively.

### Native gel mobility shift assays

Native-PAGE was used to assess the shift in mobilities of the RecBCD complex in the presence of DNA and gp5.9. For Figure 1e, the 10 μL reactions contained 25 mM Tris pH 7.5, 10 mM NaCl, 6 mM MgCl_2_. 50 nM of either RecBCD or RecBCD-Abc2 complex were mixed with 1 μM gp5.9 (or buffer), incubated for 5 minutes at room temperature and then added to 2.5 nM of a 5’-Cy5-labelled 25bp duplex DNA substrate. Reactions were then loaded onto a 6% (w/v) native polyacrylamide gel (1xTBE), then run in 1x TBE running buffer for 35 mins at a constant voltage of 150V. The gels were imaged using a Typhoon, scanning for the Cy5 fluorophore. For Figure 1f, the 20 μL reactions contained 25 mM Tris pH 7.5, 10 mM NaCl, 6 mM MgCl_2_ and 10% glycerol. 400 nM RecBCD or RecBCD-Abc2 were mixed with 4 μM gp5.9 (or buffer) and incubated for 5 mins prior to addition of 500 nM of 5’-Cy5-labelled 25bp duplex DNA substrate. The reactions were incubated for a further 5 mins at room temperature prior loading onto a 6% (w/v) native polyacrylamide gel (1xTBE). The gel was run in 1x TBE running buffer for 100 mins at a constant voltage of 150V. The gels were visualised using the Typhoon to scan for the Cy5 fluorophore (to visualise DNA) and then after stained with Coomassie blue to visualise the proteins.

**Figure 1.**
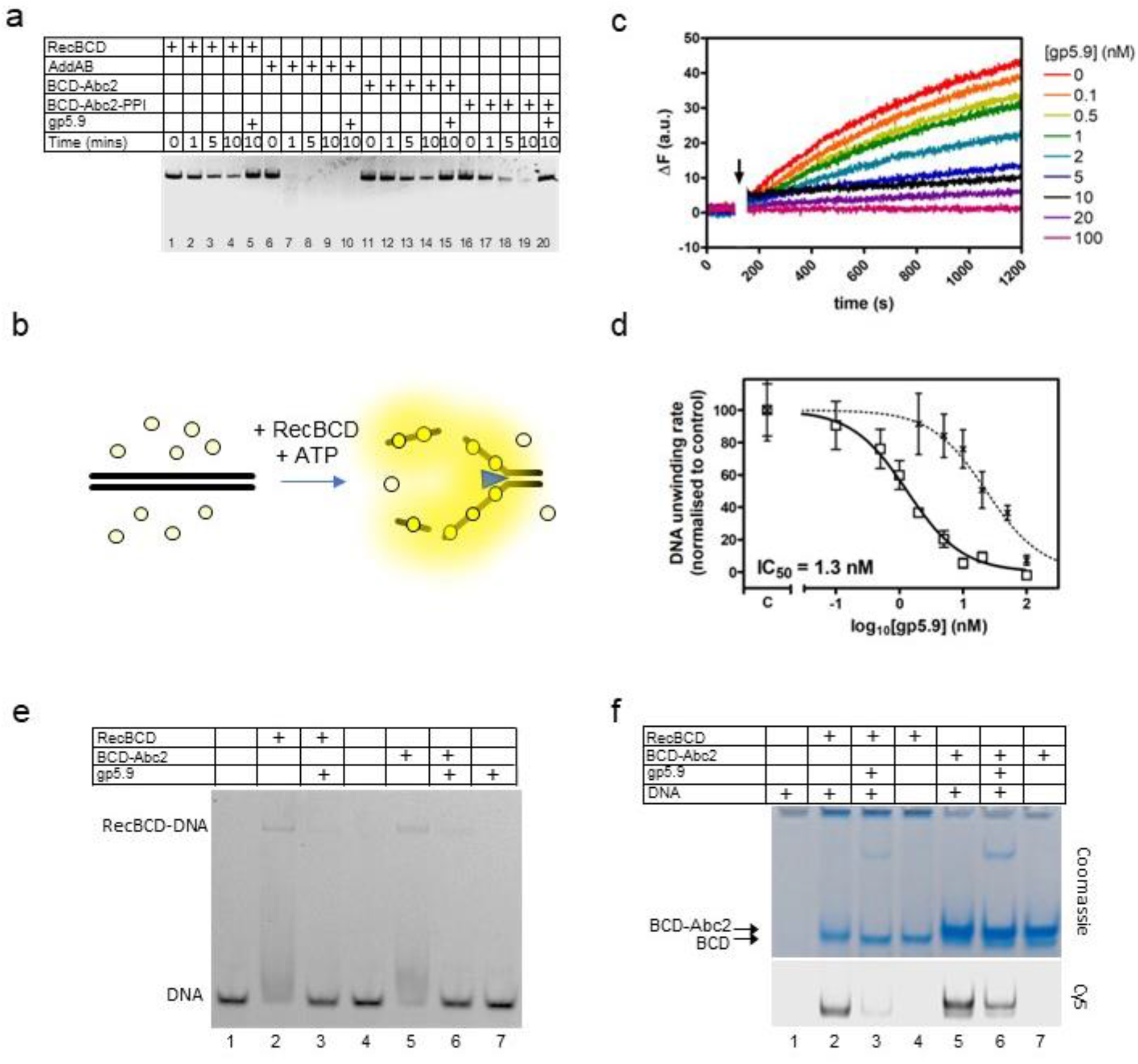
RecBCD helicase activity is inhibited by gp5.9 but not Abc2. (a) Gel-based DSB resection assay. The indicated proteins or protein complexes were incubated with a linearised dsDNA substrate for the indicated times in the presence of ATP. RecBCD (or AddAB) activity leads to DNA strand separation and depletion of the substrate. Substrate depletion is not affected by the presence of Abc2 or PPI. The substrate is not unwound by RecBCD in the presence of gp5.9 regardless of the presence of Abc2/PPI, but gp5.9 does not inhibit the orthologous AddAB complex. (b) Schematic of the DNA helicase assay using the fSSB biosensor (yellow circles). The ssDNA products of DNA unwinding are bound by fSSB causing a fluorescence increase. (c) Representative example of DNA unwinding traces for RecBCD in the presence of increasing [gp5.9]. The black arrow indicates addition of ATP to initiate the reaction. (d) A plot of initial DNA unwinding rate versus [gp5.9] yields a low nanomolar inhibition constant (IC_50_). The two data sets shown are for gp5.9 with the N-terminal his-tag removed (squares and solid fit line; IC_50_ = 1.3 nM) and with the his-tag intact (crosses and dotted fit line; IC_50_ = 23 nM). Values for the unwinding rate are normalised to a control experiment in the absence of gp5.9 (marked C on the x-axis). (e) Electrophoretic mobility shift assay showing the effect of gp5.9 and Abc2 on the DNA binding activity of RecBCD. A Cy5-labelled blunt-ended duplex DNA substate was incubated with the proteins indicated and then run on a native gel to separate the free and bound DNA species. (f) Native “inverse EMSA” assay showing binding of DNA by RecBCD or RecBCD-Abc2 complexes, and the effect of gp5.9 on both systems. The indicated complexes were incubated with or without labelled DNA substrate, either in the presence or absence of gp5.9 as indicated. The samples were then run on native polyacrylamide gels and stained with Coomassie to reveal the position of the intact protein complexes. The same gel was also imaged for Cy5 fluorescence to reveal the position of the DNA. Note that free DNA runs off the bottom of the gel in this experiment, whereas bound DNA co-migrates with the RecBCD complexes.

### Cryo-EM grid preparation

RecBCD and gp5.9 proteins were thawed and mixed in buffer B100 on ice at final concentrations of 0.3 μM RecBCD, 0.9 μM gp5.9 (based on monomeric weight) and 5 mM MgCl_2_. Quantifoil-Au 2/1 μm holey carbon film grids (300 mesh) were cleaned by successive washes of MilliQ water and ethylacetate. They were then washed with 0.3 mM n-dodecyl-beta-D-maltoside (DDM) before being immediately covered with a solution of diluted graphene oxide sheets in 0.3 mM DDM, as described previously (5). A sample volume of 4 μL was evenly applied to the graphene oxide coated carbon-side of the grid and frozen in liquid ethane using a Vitrobot Mark IV (FEI) with a 10 s wait time and 1.5 s blot time respectively. The Vitrobot chamber was maintained at close to 100% humidity and 4°C. RecB^D^CD-Abc2 was thawed and mixed with a 1.5 fold molar excess of the hairpin DNA substrate for 10 min at room temperature before being placed on ice and ligands and buffer B100 added to give the desired concentrations. The final mixture contained 0.2 μM RecB^D^CD-Abc2, 0.3 μM DNA, 5 mM magnesium chloride and 2 mM ADPNP. C-flat 2/1 μm holey carbon film grids (400 mesh) were covered with a solution of diluted graphene oxide sheets in 0.3 mM DDM, as described previously (5). The sample (3 μL) was evenly applied to the graphene oxide coated side of the grid and frozen in liquid ethane using a Vitrobot Mark IV (FEI) with a 2 s wait time and 1.5 s blot time respectively. The Vitrobot chamber was maintained at close to 100% humidity and 4°C.

### Cryo-EM data collection

The RecBCD-gp5.9 cryo-EM dataset was collected at LonCEM (The Francis Crick Institute, London) using a Titan Krios microscope operated at 300 kV with a Gatan K3 detector in super resolution mode. A nominal magnification of 81,000x was set yielding a pixel size of 1.10 Å. A total of 5,064 images were collected with a nominal defocus range of −1.0 to −2.5 μm in 0.3 μm increments. Each image consisted of a movie stack of 40 frames with a total dose of ~50 e-/Å^2^ over 4.3 s, corresponding to a dose rate of ~14 e^−^/pixel/s.

The RecB^D^CD-Abc2-DNA-ADPNP cryo-EM dataset was collected at eBIC (Diamond Light Source, UK) using a Titan Krios microscope operated at 300 kV with a Gatan K2 detector in counting mode. A nominal magnification of 130,000 was set, yielding a pixel size of 1.06 Å. A total of 2,500 images were collected with a nominal defocus range of −1.2 to −2.4 μm in 0.3 μm increments. Each image consisted of a movie stack of 30 frames with a total dose of ~56 e-/Å^2^ over 12 s, corresponding to a dose rate of ~5.2 e^−^/pixel/s.

### Cryo-EM data processing – gp5.9

The 5,064 movie stacks were aligned and summed using Motioncor2 (25) before CTF parameters were estimated for each micrograph using Gctf (26). Outlying poor quality images, based on CTF figure of merit, defocus value and predicted resolution were removed to give 4,080 micrographs for further processing. Template-based particle picking was done with Gautomatch, using 25 Å lowpass-filtered (LPF) 2D-reprojections of an 8 Å resolution RecBCD-gp5.9 cryo-EM map obtained from in-house data on a Technai F20 microscope operating with a Falcon2 detector. A total of 1,458,124 picked particles were extracted 2x binned in RELION3 (27). To remove picking artefacts and noise, two successive rounds of 2D classification were carried out in cryosparc2 (28) from which 674,546 potential RecBCD particles were kept (**Supplementary Figure 2b**). A further round of cleaning was carried out using template-free ab-initio classification from which two featureless classes were removed to leave 547,993 particles which represented a RecBCD-gp5.9 complex (**Supplementary Figure 2c**). These were refined in cryosparc2, using the ab initio map low-pass filtered to 20 Å as a template, to centre the particles prior to re-extraction without binning in RELION3.

The unbinned, centred particles were refined in RELION3, with the template low-pass filtered to 20 Å and with a soft, extended, 16 Å low-pass filtered mask around the complex, producing a map at a resolution of 3.68 Å (gold-standard, FSC=0.143, as with all succeeding resolution estimates) after postprocessing. The output particles were then subjected to Bayesian polishing and refined as before in RELION3 producing a map at a resolution of 3.52 Å. A round of CTF refinement of the per-particle defocus values significantly improved the map, which was subsequently refined to a resolution of 3.08 Å (**Supplementary Figure 2d**). A round of 3D classification was then with local angular searches and without a mask to separate any heterogeneity within the complex into three classes (**Supplementary Figure 2e**). A class containing 163,484 particles (30% of those classified) with strong density for the full complex was selected out from low-resolution classes with mixed occupancies and refined to a resolution of 3.20 Å (**Supplementary Figure 2f**). A focused 3D classification was then run using a soft mask around the gp5.9 and RecB arm domain region of the complex and without image alignment. This led to the exclusion of 22,026 particles with no gp5.9 density and selection of 141,458 particles with strong gp5.9 density (**Supplementary Figure 2g**), which were refined to yield a final map at 3.2 Å (**Supplementary Figure 2h**), which was deposited with a sharpening factor of −50 e^−^/Å^2^ applied. Due to the reduced resolution still observed around the C-terminus of gp5.9 and the RecB arm in the map, the refinement was continued using the local mask from the previous classification to generate a secondary map to aid model building and interpretation in this region.

### Cryo-EM data processing – Abc2

The movie stacks were aligned and summed using Motioncor2 (25) before CTF parameters were estimated for each micrograph using Gctf (26). Images with significant ice contamination were removed to give 2,338 micrographs for further processing. Template-based particle picking was done with Gautomatch, using 25 Å lowpass-filtered (LPF) 2D-reprojections of the published RecBCD-DNA-ADPNP cryo-EM map (EMD-4038, (22)). A total of 467,613 picked particles were extracted 2x binned. To remove picking artefacts and noise, two successive rounds of 2D classification were carried out in RELION3 (27) from which 185,881 potential RecBCD particles were kept (**Supplementary Figure 5b**). These were used for an unmasked 3D refinement with the EMD-4038 map filtered to 30 Å resolution as the starting template to centre the particles before re-extraction without binning. This was followed by 3D refinement with a soft mask around the full complex, Bayesian polishing and CTF refinement of the per-particle defocus values. The map resolution improved from 3.6 Å (0.143 FSC cutoff, RELION3) to 3.3 Å after Bayesian polishing, CTF refinement and a further 2D classification without image alignment to select 167,080 highly ordered particles (**Supplementary Figure 5c and 5d**).

To separate different states/occupancies of the complex, a focused refinement was done on the polished particles with a soft mask around Abc2 and the surrounding domains of RecC, followed by 3D classification without image alignment using the same focused mask (**Supplementary Figure 5e and 5f**). The majority of the particles (71% of those classified) formed a high-resolution class with strong density for the full complex, including Abc2 bound on the surface of RecC. A minor, low-resolution class was excluded containing 7% of the particles respectively. The remaining particles (22 % of those classified) were in a class that displayed additional density attached to Abc2 for the contaminant *E.coli* PpiB protein, representing the RecBCD-DNA-Abc2-PPI complex. The 119,163 RecBCD-DNA-Abc2 particles were refined to generate a map at a resolution of 3.4 Å (0.143 FSC cutoff, RELION3) deposited with a sharpening B-factor of −50 Å^2^ (**Supplementary Figure 5g and 5i, Table 1**.). The 37,072 RecBCD-DNA-Abc2-PPI particles were refined to yield a map at a resolution of 3.8 Å (0.143 FSC cutoff, RELION3), which was deposited with a sharpening B-factor of −50 Å^2^ (**Supplementary Figure 5h and 5i, Table 1**).

**Table 1.** Cryo-EM data collection, refinement and validation statistics.

|  | gp5.9-RecBCD<br>(EMDB-15803)<br>(PDB 8B1R) | Abc2-RecB<sup>D</sup>CD-DNA-ADPNP<br>(EMDB-15804)<br>(PDB 8B1T) | Abc2-RecB<sup>D</sup>CD-PpiB-DNA-ADPNP<br>(EMDB-15805)<br>(PDB 8B1U) |
| --- | --- | --- | --- |
| **Data collection and processing** |  |  |  |
| Magnification | 81,000 | 130,000 |  |
| Voltage (kV) | 300 | 300 |  |
| Detector | K3 | K2 |  |
| Energy slit width (eV) | 20 | 20 |  |
| Electron exposure (e<sup>-</sup>/Å<sup>2</sup>) | 50 | 56 |  |
| Exposure rate (e<sup>-</sup>/pixel/s) | 14.0 | 5.2 |  |
| Defocus range (μm) | -1.0 to -2.5 | -1.2 to -2.4 |  |
| Pixel size (Å) | 1.10 | 1.06 |  |
| Movies collected | 5,064 | 2,500 |  |
| Initial particle images (no.) | 674,546 | 185,881 |  |
| Final particle images (no.) | 141,458 | 119,163 | 37,072 |
| Symmetry imposed | C1 | C1 | C1 |
| Map resolution (Å) | 3.2 | 3.4 | 3.8 |
| FSC threshold | 0.143 | 0.143 | 0.143 |
| Map resolution range (Å) |  |  |  |
| **Refinement** |  |  |  |
| Initial model used (PDB code) | 5MBV | 5LD2, 8B1R | 8B1T |
| Map sharpening *B* factor (Å<sup>2</sup>) | -50 | -50 | -50 |
| Model resolution (Å) | 3.1 | 3.3 | 3.7 |
| FSC threshold | 0.143 | 0.143 | 0.143 |
| Model to map correlation | 0.87 | 0.83 | 0.78 |
| Model composition |  |  |  |
| Non-hydrogen atoms | 21,953 | 23,986 | 24,127 |
| Protein residues | 2,746 | 2,869 | 2,886 |
| DNA residues | - | 51 | 51 |
| Ligand molecules | 1 (Mg<sup>2+</sup>) | 1 (ADPNP)<br>1 (Mg<sup>2+</sup>) | 1 (ADPNP)<br>1 (Mg<sup>2+</sup>) |
| *B* factors (Å<sup>2</sup>) |  |  |  |
| Protein | 89.5 | 72.0 | 68.0 |
| DNA | - | 168.5 | 162.6 |
| Ligand | 61.0 | 50.0 | 54.5 |
| R.m.s. deviations |  |  |  |
| Bond lengths (Å) | 0.004 | 0.003 | 0.002 |
| Bond angles (°) | 0.617 | 0.493 | 0.464 |
| Validation |  |  |  |
| MolProbity score | 1.4 | 1.5 | 1.5 |
| Clashscore | 7.5 | 6.0 | 5.7 |
| Poor rotamers (%) | 0.5 | 1.4 | 1.5 |
| Ramachandran plot |  |  |  |
| Favored (%) | 99.3 | 98.9 | 98.9 |
| Allowed (%) | 0.7 | 1.1 | 1.1 |
| Disallowed (%) | 0.0 | 0.0 | 0.0 |

### Model building and refinement

For the 3.2 Å resolution RecBCD-gp5.9 map, RecBCD from the published RecBCD-GamS structure (PDB ID: 5MBV, (9)) was docked into the density using Chimera (29), and refined with jelly-body restraints using Refmac5 (29) in CCPEM (30). A focused refinement in RELION3 with the mask around the gp5.9 binding site used during data processing (Supplementary Figure 2g) generated a more resolved map to facilitate the building of gp5.9 and remodelling of interacting regions of the RecB arm and RecC C-terminal domains respectively in coot (32). Initially a long idealised α-helix was docked in to start gp5.9 building with an N-terminal β-strand extension clearly visible. At this resolution and with the majority of the peptide ordered, sequence assignment was clear and straightforward with residues 1-49 modelled. The chain was duplicated and docked into the density for the second gp5.9 molecule within the homo-dimer, with only minor adjustments required for the very N-terminal residues and some altered side chain conformers (Supplementary Figure 3). An alphafold (33) model of gp5.9 was later used to help validate our model. Notably, an alphafold model of the RecBCD complex combined with observations of inconsistencies between the model and map in the RecB arm led to the finding that the protein register within this domain was off by seven residues for residues 241-289, which are flanked by disordered loops in prior RecBCD structures. Two tryptophan residues seven residues a part in this region had become the basis for the misalignment in the first model built for RecBCD (24), and it was not uncovered with the ambiguous nature of the density in this region in the subsequent RecBCD structures that followed, until this significantly higher resolution gp5.9 complex map (**Supplementary Figure 11**). We went back through past RecBCD complex structures and found that the new RecB arm model fit significantly better into all of the past maps. Comparative analyses of RecBCD structures in this paper use the corrected models.

The published RecBCD-DNA-ADPNP structure (PDB ID: 5LD2, (22)) with the corrected RecB arm domain, was docked into the 3.4 Å Abc2-containing map using Chimera and refined with jelly-body restraints using Refmac5(29) in CCPEM. Additional density was present on the surface of RecC that was sufficiently ordered and resolved to facilitate manual building of the novel Abc2 polypeptide chain from residues 6-52 (**Supplementary Figure 8a**) in Coot with iterative rounds of manual building and Phenix real-space refinement (31). Initially, three α-helices were docked in and connecting loops built. Sequence was assigned based on a combination of secondary structure prediction generated by JPRED (32) and register matching of density size and environment to the chemical properties of the known amino acid sequence. Concurrently, the surrounding RecC region was adjusted to fit the density with minor remodelling except for a surface bundle including residues 252-294 (**Supplementary Figure 7b**). This helical bundle peels back to allow Abc2 binding and then loosely clamps back to hold Abc2 in place, although the density in its new position was insufficient for model building (**Supplementary Figure 7c**).

For the Abc2 and PPI-containing map at 3.8 Å resolution, the final RecBCD-DNA-Abc2 model was docked in and refined with jelly-body restraints as described above. As well as containing additional density for PPI, the map showed further density for building an extended part of the C-terminal region of the Abc2 model up to residue 66 (**Supplementary Figure 8b and 9a**). The density in this region was ambiguous as indicated by local resolution estimation identifying the resolution to be worse than 5 Å for PpiB (**Supplementary Figure 5h**). As a result, the crystal structure of *E.coli* PpiB (PDB: 1LOP) was docked into the density (**Supplementary Figure 8**) for the purpose of figures but not for the deposition of a 3D model. It is unclear what changes are induced in Abc2 and in PPI itself upon PPI binding to the C-terminal portion of Abc2. However, the model is strongly suggestive of P68 of Abc2 binding within the active site of the prolyl isomerase (**Figure 4f**). The final models of each structure were real-space refined in Phenix against the deposited maps with ADP-factor refinement enabled and model quality assessed by MolProbity (33) to generate the final model statistics (**Table 1**).

## Results

### Gp5.9, but not Abc2, inhibits RecBCD helicase activity

The T7 gp5.9 protein was shown previously to inhibit RecBCD but the mechanism was not determined. Therefore, we made recombinant gp5.9 and studied its effects on RecBCD activity *in vitro*. We found that gp5.9 expression was toxic to *E. coli*, and therefore used insect cell expression to express and purify the protein with a cleavable histidine tag (**Supplementary Figure 1**). Unless stated otherwise, all experiments were performed with a preparation of gp5.9 with the tag removed. We first assessed the effect of T7 gp5.9 on RecBCD helicase-nuclease activity using a simple gel-based DSB resection assay. As expected, RecBCD rapidly unwound a linearised duplex DNA substrate (**Figure 1a,** lanes 1-4). In the presence of excess purified gp5.9 (1 μM), the RecBCD enzyme was strongly inhibited (compare lanes 4 and 5). To assess the specificity of gp5.9 for RecBCD, we performed analogous experiments using the AddAB helicase-nuclease, a RecBCD orthologue from *Bacillus subtilis* (34), and observed no inhibition of DNA degradation (lanes 6-10). This strongly suggests that gp5.9 acts by binding directly and specifically to the RecBCD complex, as opposed to blocking DNA ends.

We next compared the effects of Abc2 on RecBCD activity. It has been established previously that Abc2 binds directly to RecBCD, but it seems to have at most subtle effects on its spectrum of activities *in vitro* (16). We were unable to purify the Abc2 polypeptide alone. Therefore, to confirm these observations, we purified the intact RecBCD-Abc2 complex which was found to co-purify with host peptidyl-prolyl cis-trans isomerase B (*E. coli* PpiB) as observed previously (14, 15) (**Supplementary Figure 1**). It was found that expression of the Pro68Ala mutant form of Abc2 (Abc2^P68A^) with RecBCD prevented co-purification with the host PpiB protein, allowing us to also isolate the RecBCD-Abc2^P68A^ complex (**Supplementary Figure 1**). Note therefore that, in the following experiments, we are comparing the biochemical activities of independently-purified samples of free RecBCD, RecBCD-Abc2^P68A^ and the RecBCD-Abc2-PPI complex. As expected based on previous work, the observed DNA unwinding activities of the three preparations were very similar (**Figure 1a,** compare lanes 1-4 with 11-14 and 16-19), and all three preparations were completely inhibited by excess gp5.9 (lanes 5, 15 and 20).

To provide more quantitative insights into RecBCD inhibition by gp5.9, we next employed a continuous multiple turnover helicase assay which uses a fluorescent biosensor for ssDNA (fSSB) to monitor DNA unwinding in real-time (23, 35, 36) (**Figure 1b**). RecBCD was pre-incubated at a low concentration (10 pM) with a linear DNA substrate (100 pM molecules) that was devoid of Chi sequences. DNA unwinding was then initiated by the addition of ATP in the presence of fSSB. A rapid increase in fluorescence indicated that RecBCD unwound the DNA over several hundred seconds (**Figure 1c**). We next repeated the experiment in the presence of increasing concentrations of gp5.9, measuring the initial rate of unwinding in each case. Increasing concentrations of gp5.9 inhibited RecBCD helicase and a plot of [gp5.9] versus unwinding rate was well-fit to an inhibitor dose-response curve yielding IC_50_ = 1.3 nM (solid line; **Figure 1d**). Equivalent experiments using a gp5.9 preparation retaining the histidine tag yielded a significantly higher IC_50_ value of 23 nM (dotted line; **Figure 1d**). This suggests that the N-terminal region of the protein is important functionally. Together, these experiments show that both gp5.9 and Abc2 bind directly to RecBCD, but only gp5.9 acts as an inhibitor of RecBCD DNA unwinding activity.

### Gp5.9 inhibits RecBCD-DNA interaction independently of Abc2 binding

To determine the mechanistic basis for inhibition of RecBCD by gp5.9, and to assess if the phage proteins could bind simultaneously to RecBCD we next performed EMSA and “inverse EMSA” experiments to monitor DNA binding by different RecBCD-inhibitor complexes. In conventional EMSA experiments a Cy5-labelled DNA substrate (25mer blunt duplex) at low concentration (5 nM) was incubated with RecBCD or RecBCD-Abc2, either in the presence or absence of excess gp5.9 (1 μM). RecBCD caused a shift in the mobility of the substrate indicative of the expected RecBCD-DNA interaction (**Figure 1e**). This gel shift was largely eliminated by the addition of gp5.9 to RecBCD before DNA was added. The RecBCD-Abc2 complex behaved identically to wild type. It was able to bind the duplex DNA substrate, but binding was largely inhibited by excess gp5.9. The phage protein alone did not interact with DNA. In complementary “inverse” EMSA experiments (**Figure 1f**), RecBCD-DNA or RecBCD-Abc2-DNA complexes were run at high concentrations in native polyacrylamide gels that were imaged for Cy5-DNA using a confocal scanner and then stained with Coomasie to detect protein-containing complexes. Note in these experiments that free DNA is not detected as it has run off the bottom of the gel (lane 1). RecBCD was able to bind to DNA as expected (lane 2). However, addition of gp5.9 before DNA dramatically reduced the amount of DNA detected and caused a small increase in the mobility of the RecBCD complex showing it was interacting directly (compare lanes 2 and 3). The RecBCD-Abc2 complex displayed a reduced mobility compared to RecBCD alone (compare lanes 2-4 with 5-7). There was evidence for a small amount of free RecBCD in the preparation in the form of a fine band running with identical mobility to the RecBCD-alone preparation. Importantly, both bands co-migrated with Cy5-DNA showing that DNA and Abc2 can bind simultaneously to RecBCD (lane 5). Pre-addition of gp5.9 to the RecBCD-Abc2 preparation reduced Cy5-DNA binding to both protein complexes (compare lanes 5 and 6). Together, these data show that the gp5.9 binding site on RecBCD is distinct from that of Abc2, and that gp5.9 inhibits DNA binding, potentially by direct competition at the DNA binding locus.

### gp5.9 is a DNA mimic protein with an unprecedented architecture

We next purified the RecBCD-gp5.9 complex (using the tag-free form of gp5.9) and analysed its structure using single particle cryoEM. The dataset was found to be relatively homogeneous with evident additional density for the phage protein on the surface of the complex (**Supplementary Figure 2**). There was structural heterogeneity only within the RecD protein, suggesting that the RecD conformation becomes uncoupled to the rest of the complex when bound to gp5.9. This was improved by 3D classification, after which an ordered high-resolution class (30% of the classified particles) was separated for the RecBCD-gp5.9 complex with the RecD 1A and 1B domains resolved but not the 2A nor SH3 domains (**Supplementary Figure 2e**). A 3.2 Å resolution cryoEM map was obtained (**Figure 2a**, **Supplementary Figure 2h**) facilitating the building of a model for two chains of gp5.9, both containing residues 1-49 (**Figure 2b** and **Supplementary Figure 3**). The structure reveals that gp5.9 adopts a parallel coiled-coil architecture in which the N-termini are slightly braced apart by a short anti-parallel beta-sheet. Many aspartate and glutamate residues are displayed on the outer surface of this exceptionally negatively-charged protein (pI = 4.0; **Figures 2b and 2c**). The phage protein engages RecBCD in a position which overlaps extensively with the DNA binding site, and also with the binding site for the DNA mimic protein Gam (**Figures 2d-f**). The C-terminal end of the coiled-coil binds to the extended RecB arm domain, while the N-terminus binds closer to the RecBCD core, making contacts with both the RecB and RecC subunits and helping to explain the reduced efficacy of inhibition by purified gp5.9 retaining an N-terminal tag (**Figure 1d**). The overall structures of RecBCD when bound to either DNA or gp5.9 are almost identical, although small rigid body domain movements help accommodate differences in the dimensions of the two ligands (**Supplementary Figure 4**).

**Figure 2.**
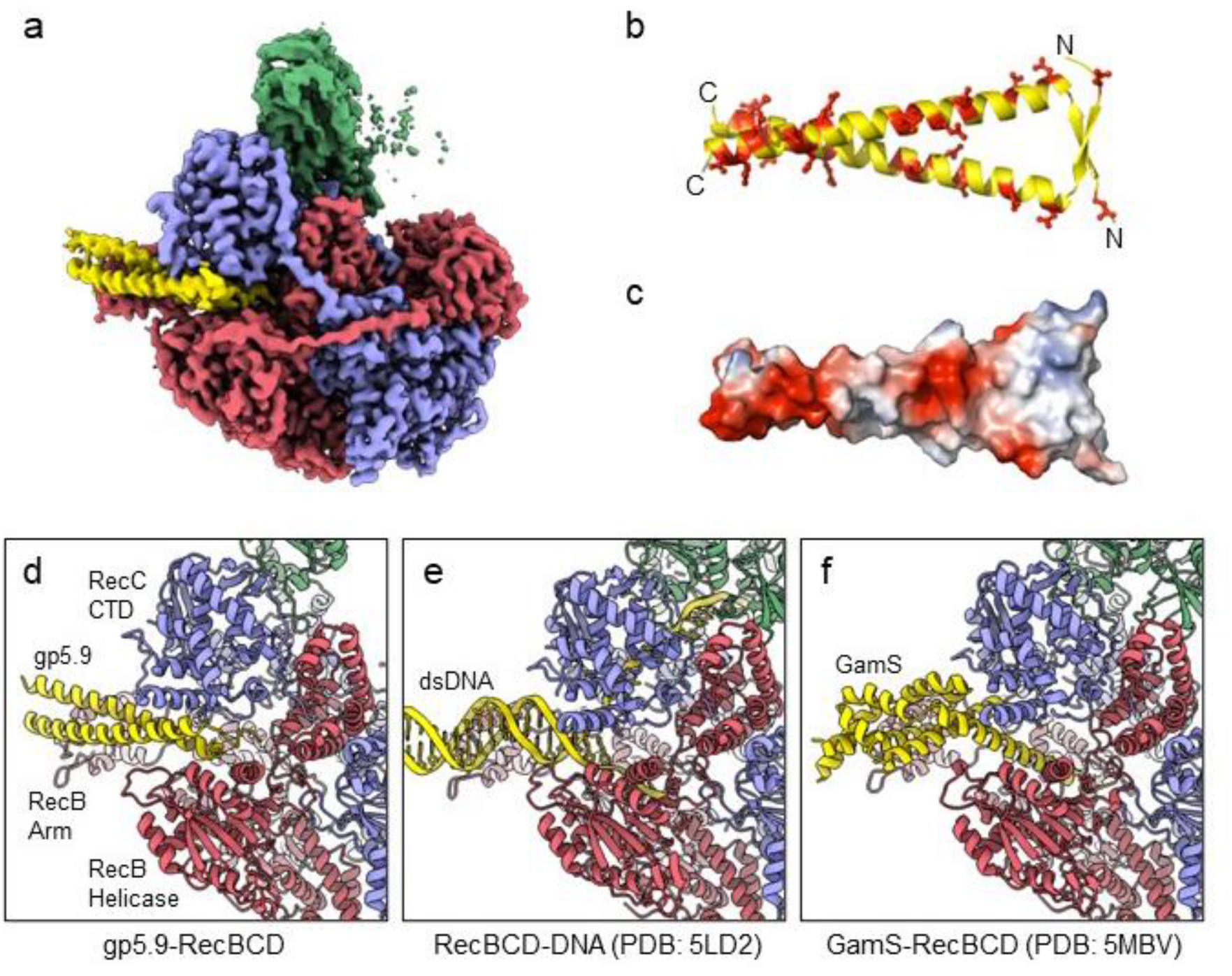
CryoEM structure of the RecBCD-gp5.9 complex. (a) CryoEM map of the RecBCD-gp5.9 complex. Subunit colour-coding is as follows: RecB in red, RecC in slate blue, RecD in green and gp5.9 in yellow. (b-c) The gp5.9 dimer adopts a parallel coiled-coil architecture braced by an N-terminal beta-sheet and displays many negatively charged Asp and Glu residues (red) on its surface. (d) Section of the atomic model of the RecBCD-gp5.9 complex with colouring as in (a). (e-f) Sections of the models of the RecBCD-DNA (PDB:5LD2) and RecBCD-Gam (PDB:5MBV) complexes for comparison.

Many ion pair contacts are formed between D/E and K/R residues in gp5.9 and RecBCD respectively (**Figure 3 and Table 2**). These imitate a subset of the contacts formed between the negatively charged DNA phosphates and K/R residues in the RecBCD-DNA complex and, on this basis, we can conclude that gp5.9 acts by DNA mimicry. Interestingly, the contacts formed between RecBCD and Gam (another DNA mimic protein) are equivalent to a different subset of the RecBCD-DNA interactions such that only a few of the charge-based interactions (with RecB Arg residues 254, 255 and 761) are conserved across all three complexes (**Figure 3 and Table 2**). Indeed, besides their dimeric, α-helical, and highly charged structures there is little else in common between the two phage proteins and the coiled-coil architecture of gp5.9 is unprecedented for a DNA mimic. In addition to charge-based interactions that mimic contacts with DNA phosphates, gp5.9 makes additional protein-protein interactions which presumably provide specificity for RecBCD, as opposed to other DNA binding proteins including AddAB.

**Figure 3.**
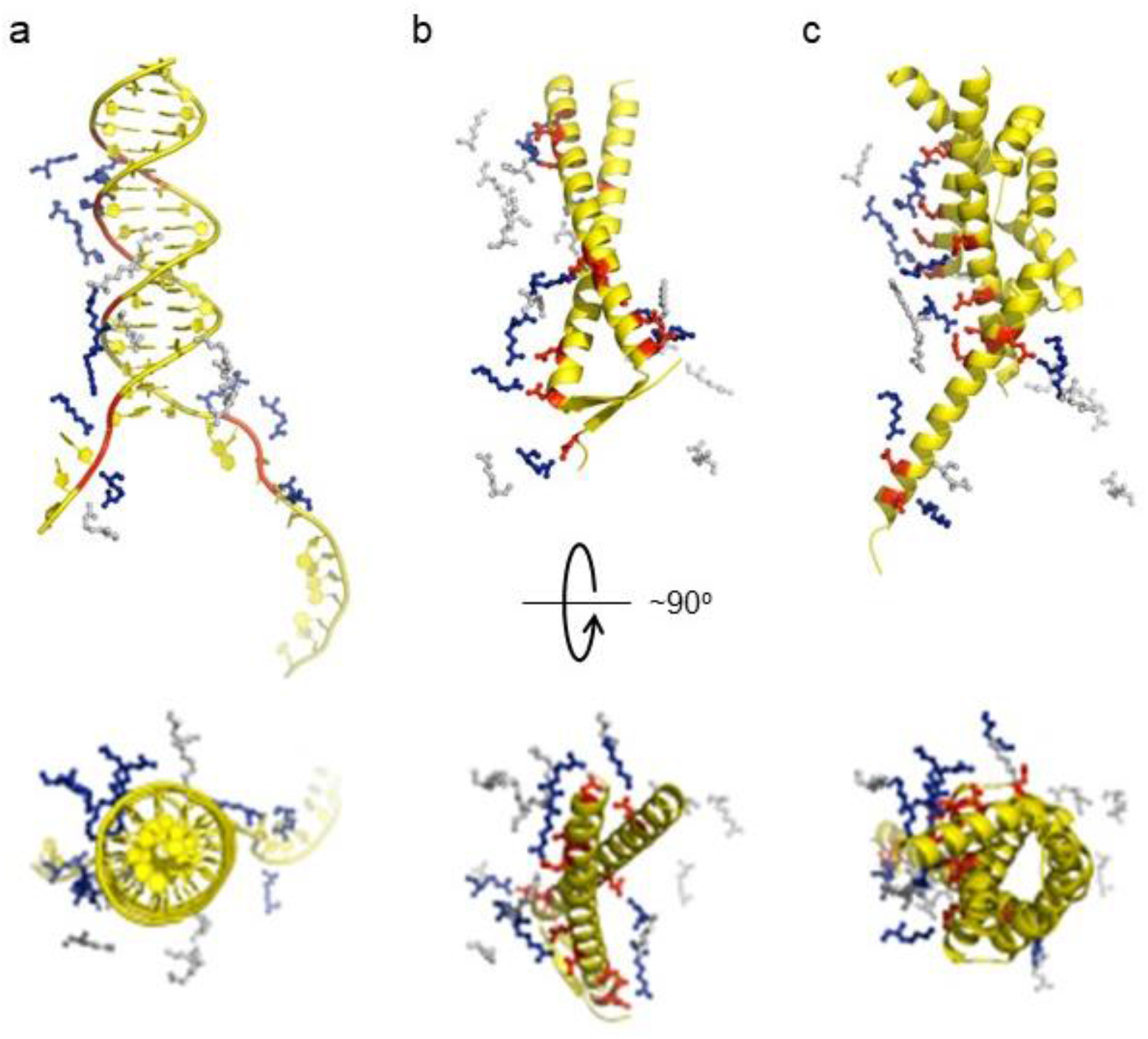
gp5.9 is a DNA mimic protein. Charge pair interactions between R/K residues in RecBCD (red) and either phosphates or D/E residues (blue) for DNA (a), gp5.9 (b) or Gam (c). The images are all taken from the same point of view for complexes that were superimposed using the RecB arm domain. The R/K residues displayed are all of those shown in **Table 2** and are the same for all three complexes. They are coloured blue if they make contacts with the ligand in each case (either DNA, gp5.9, or Gam) and grey if they do not.

**Table 2.** Gp5.9 is a DNA mimic protein. Ion pair contacts (<4Å) between Arg/Lys residues in the RecBCD complex and negatively charged moieties in either DNA (phosphates) or DNA mimic proteins (Asp/Glu). Shaded rows highlight charge-charge interactions that are analogous across all three complexes. These interactions are illustrated in **Figure 3**.

| **RecBCD**<br>(subunit indicated) | **DNA** | **Gp5.9** | **Gam** |
| --- | --- | --- | --- |
| R254 (B) | O3'-25 | E45 | D73 |
| R255 (B) | O3'-56 | D38 / E39 | E118 / E125 |
| K256 (B) | OP1-57 / O5'-57 |  | E118 |
| K264 (B) |  | E36 |  |
| K268 (B) |  |  | D107 |
| K288 (B) | OP-59 |  | E111 |
| R297 (B) | OP-58 |  | E114 |
| K299 (B) | OP-27 |  |  |
| R561 (B) | OP-69 | D4 |  |
| R584 (B) |  |  | D48 |
| R761 (B) | O5'-68 / O3'-67 | D11 | E51 |
| R822 (B) | OP-67 | D15 |  |
| R823 (B) | OP-18 |  |  |
| R824 (B) |  | D21 / E24 | E65 / E68 / E125 |
| K828 (B) |  |  | E118 |
| R846 (C) | OP-10 / O3'-9 |  |  |
| R968 (C) | OP-11 / O3'-10 |  |  |
| R1001 (C) | OP-12 |  |  |
| K1066 (C) |  |  | E70 / D74 |
| R1068 (C) |  | D11/D15 |  |
| K1070 (C) |  | E24 | E70 |

### Abc2 binds to the RecC subunit, close to a putative exit channel for recombinogenic DNA

We next solved cryoEM structures of the RecBCD-Abc2-PPI complex bound to a tailed DNA substrate (**Figure 4**). All of the particles within the dataset contained ordered density for RecBCD with additional density evident for the Abc2 protein (**Supplementary Figure 5**). Focused classification with a mask around the phage binding site (**Supplementary Figure 5f**) isolated 71% of the particles into a homogeneous class from which a 3.4 Å resolution structure of the RecBCD-Abc2-DNA complex was solved (**Figure 4a, Supplementary Figure 5g**). A second class containing 22% of the data displayed further additional resolved density in the vicinity of Abc2, corresponding to the *E.coli* PpiB protein. From this class, a 3.8 Å resolution structure of RecBCD-Abc2-PPI-DNA complex was solved (**Figure 4b, Supplementary Figure 5h**). The Abc2 protein adopts an extended alpha-helical structure and binds to the surface of the “inactivated helicase domains” of RecC which play a key role in Chi recognition (**Figure 4c-4e**) (24). Consistent with biochemical observations, this location suggests a role for Abc2 in modulating the late stages of the DSB processing reaction catalysed by RecBCD (i.e. Chi recognition and associated conformational changes mediated by a “latch” structure in RecC, or the subsequent loading of RecA protein). The binding site is also close to the interface with the helicase domains of RecB, and to a tunnel between RecB and RecC (**Supplementary Figure 6)** which has been suggested to allow exit of a recombinogenic ssDNA loop (the site of RecA loading) after Chi recognition (5, 37). In the RecBCD-Abc2-PPI structure, RecBCD and Abc2 are essentially identical, but Abc2 is also bound to the host PpiB protein. Abc2 interacts extensively with the Chi-recognition domains of RecC as before, but there are no interactions between PpiB and RecBCD (**Figure 4c**). The structure can be superposed onto the corresponding translocation activated RecBCD-DNA structure (PDB: 5LD2; (21)), containing the same DNA substrate and non-hydrolysable ATP analogue, ADPNP (**Supplementary Figure 7a**). This shows that Abc2 has no significant effect on the conformation of the majority of the RecBCD complex except for a minor opening of the nuclease domain relative to RecC and RecD. Additionally, Abc2 binding displaces a helical bundle, residues 252-294, from the RecC surface (**Supplementary Figure 7b**) and some partial density can be seen for the displaced region clamping back over the exposed face of Abc2 (**Supplementary Figure 7c**).

**Figure 4.**
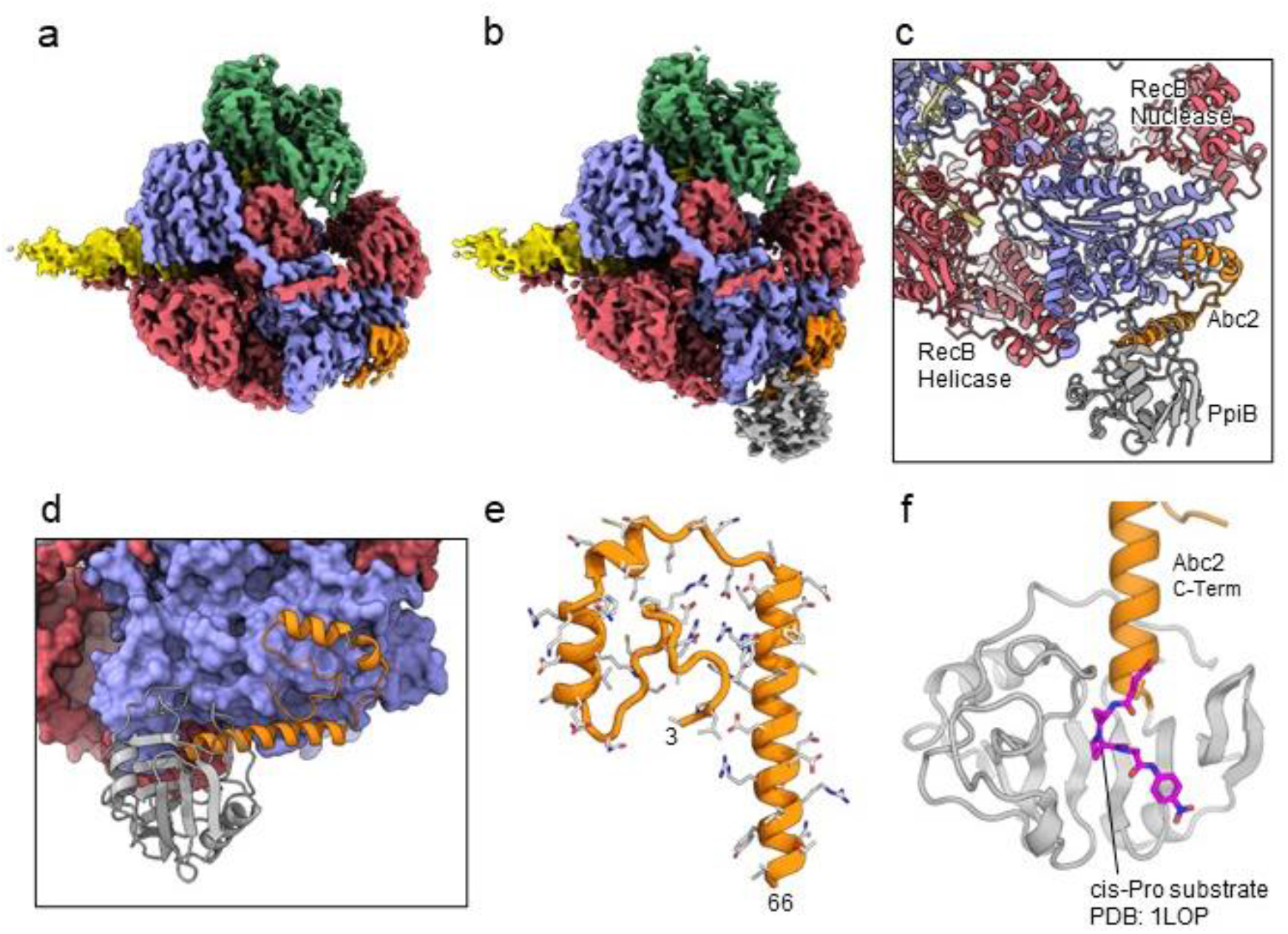
CryoEM structures of the RecBCD-Abc2 and RecBCD-Abc2-PPI complexes. (a) CryoEM map of the RecBCD-Abc2-DNA complex, coloured as in Figure 2 but with the DNA substrate yellow and Abc2 in orange (b) CryoEM map of the RecBCD-Abc2-PpiB-DNA complex, with PpiB in grey. (c) Section of the atomic model of the RecBCD-Abc2-PpiB-DNA complex. (d) Surface view of the Abc2 binding site on the surface of the RecC Chi-binding domains. (e) Atomic model of residues 3-66 of Abc2 in the PpiB-bound complex showing an extended helical structure. (f) Model of the C-terminus of Abc2 bound to PpiB. A substrate analogue (magenta) is superimposed on the PpiB structure based upon PDB:1LOP. The structure strongly suggests that P68 (two residues beyond the final modelled amino acid 66) would reside in the cis-trans isomerisation active site.

The C-terminal region of Abc2 is disordered to differing extents in the structures; helix-2 of Abc2 is ordered up to residue 52 on the surface of RecC (**Supplementary Figure 8a**) but in the PpiB-bound state this helix is extended with Abc2 modelled up to residue 66 (**Supplementary Figure 8b**). This C-terminal extension protrudes into the PpiB density, into which a high resolution crystal structure of PpiB containing a bound substrate analogue (PDB:1LOP; (38)) can be docked (**Figure 4f**). The resulting model strongly suggests that P68 from Abc2 would occupy the *cis-trans* isomerisation active site in PpiB. A *cis* proline conformation at this position would disrupt the helical secondary structure and potentially direct the disordered C-terminus of Abc2 towards the RecB-RecC interface. In fact, when the RecBCD-Abc2-PPI map is blurred and contoured down, a continuous weak density path can be observed for the continuation of the Abc2 C-terminus through the PpiB binding cleft and out towards the RecB helicase domains (**Supplementary Figure 8c**). Looking around the RecB helicase domains, additional unaccountable patches of density are found above the noise in-between the 1A and 2A motor domains which enclose the ATP-binding site (**Supplementary Figure 9a**). This additional density is not seen in the Abc2 complex without bound PpiB (**Supplementary Figure 9b**). AlphaFold (33) models of the Abc2 C-terminus are highly divergent and cannot be used to interpret the density (**Supplementary Figure 10**).

## Discussion

In this work we used biochemical and structural methods to study the interactions of two phage proteins, T7 gp5.9 and P22 Abc2, with the *E. coli* RecBCD complex. We found that low nanomolar concentrations of gp5.9 specifically inhibit RecBCD helicase activity *in vitro* by preventing DNA substrate binding. Our structure of the RecBCD-gp5.9 complex explains the biochemical observations by showing how gp5.9 competes directly for the DNA binding site through mimicry of the natural DNA substrate.

The T7 gp5.9 protein is functionally analogous to Gam from phage λ (8). Moreover, both are small, dimeric and highly acidic proteins which form a compact and predominantly alpha-helical structure (6). Despite these similarities, their primary structures, overall folds and the molecular-level details of their interaction with RecBCD are different. Indeed, DNA mimic proteins display a remarkable structural diversity (13, 14) and, to the best of our knowledge, the coiled-coil architecture observed here for gp5.9 has not been observed in nature before this study. Interestingly however, this fold has been used previously as the structural framework for the rational design of synthetic DNA mimics targeting restriction endonucleases (39). Our work here validates the choice of coiled-coils for producing such synthetic proteins, and may help to inform their design against novel targets.

In cells containing the bacterial retron Ec48, the presence of either gp5.9 or Gam (either of which are indicative of phage infection) triggers programmed cell death as an abortive infection mechanism (3). Because the Ec48 system is not tolerated in a *ΔrecB* mutant, and because Gam was known to bind mainly to RecB, it was suggested that the retron binds directly to RecB and its release is triggered by Gam or gp5.9 to activate downstream effectors of cell suicide by unknown mechanisms. This model implies that Gam and gp5.9 bind to the same interface on RecB. Our results here are consistent with this hypothesis because Gam and gp5.9 do indeed share the same binding location on the RecB arm domain and other nearby regions of RecB and RecC. Therefore, DNA, Gam, gp 5.9 and possibly the retron all compete for the same site on RecBCD. In retron-free *Escherichia coli* cells, expression of gp5.9 likely sequesters all of the available RecBCD complex (present at only ~10 copies per cell (40–42)) in an inactive form, thereby protecting T7 DNA from degradation and prioritising the use of phage recombinases (**Figure 5**). In addition to its natural roles, gp5.9 may also find use as a biotechnology tool for enhancing recombineering efficiency, improving cell-free transcription-translation systems, or stabilising DNA nanostructures *in vivo* through its ability to potently inhibit RecBCD exonuclease activity (43–45).

**Figure 5.**
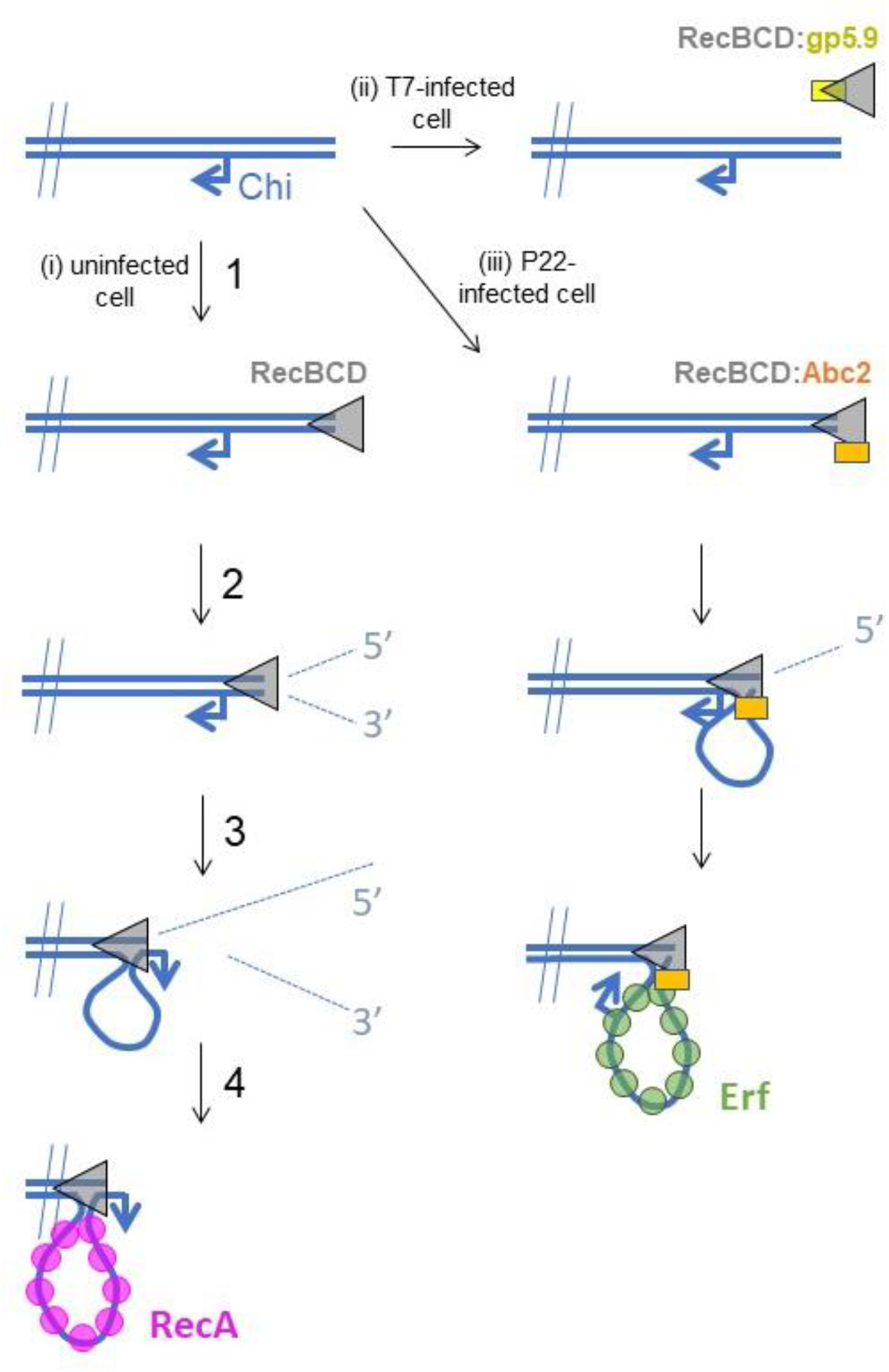
Hypothetical models for control of RecBCD by T7 gp5.9 and P22 Abc2. In uninfected cells (pathway i), free DNA ends are processed by RecBCD to yield a recombinogenic 3’-terminated ssDNA overhang coated with host RecA protein. The reaction proceeds in five steps. (1) RecBCD binds to the free DNA end. (2) RecBCD translocates and unwinds the duplex DNA degrading both strands as it progresses. This activity is detrimental to phage as it can help degrade invading linear DNA species and initiates CRISPR acquisition. (3) RecBCD recognises a Chi sequence (these are over-represented in host DNA). (4) RecBCD continues to unwind DNA and to degrade the 5’-terminated DNA strand, but the 3’-terminated strand is protected from degradation and forms an expanding loop structure. (5) RecBCD loads RecA protein onto the ssDNA loop ready for subsequent strand invasion and further steps of DNA repair. In T7-infected cells (pathway ii), the gp5.9 protein titrates RecBCD by competing for the DNA binding site. Linear phage DNA cannot be degraded by RecBCD and is available for phage recombination and replication. In P22-infected cells (pathway iii), Abc2 binds to the RecBCD complex and modifies the enzyme to load the P22 recombinase (Erf), as opposed to RecA, onto the ssDNA products. Recombinase loading might occur either constitutively as indicated here, or in response to Chi-recognition as in the unperturbed pathway.

The effects of Abc2 on RecBCD activity are complex and remain less well understood than is the case for the two DNA mimic proteins. Expression of Abc2 in *E. coli* eliminates RecBCD-dependent recombination but exonuclease activity is retained (18). Purified Abc2 has only subtle effects on RecBCD activity *in vitro* which are somewhat similar to the effects elicited upon recognition of Chi sequences (9, 15–18). Based upon their observations, Murphy and co-workers developed a model in which Abc2 hijacks RecBCD to generate the ssDNA required for phage recombination and modifies its RecA loading function to instead recruit its own recombinase (called Erf) to the ssDNA products (**Figure 5**). Our biochemical and structural data are consistent with this hypothesis. Abc2 is bound to RecC in close proximity to both a “latch helix” which is thought to control conformational changes in response to Chi, and to an “alternative exit channel” in the RecBCD complex which is thought to accommodate a ssDNA loop in the post-Chi state (i.e. the recombinogenic mode) (5, 37, 46). Under normal circumstances, this loop would be loaded with *E. coli* RecA protein to promote subsequent steps in the host DNA break repair pathway. Given its location, it is plausible that Abc2 prevents RecA loading at this site and/or promotes the binding of Erf to the emerging ssDNA, but this remains to be tested experimentally. In addition to Abc2 and Erf, the P22 recombination system also includes the uncharacterised proteins Abc1 (Anti-RecBCD protein 1) and Arf (Accessory recombination factor) which are both required for full activity during P22 infection (17). The role of Abc1 is unknown and its ability to bind directly to RecBCD has never been tested. Likewise, exactly how Arf influences P22 recombination is unclear, but its small size and acidic primary sequence hint that it might be a DNA mimic protein. Finally, the significance of the interaction between Abc2 and host PpiB is not clear. Our structure shows that the PPI active site almost certainly engages P68 of Abc2, and it is therefore plausible that proline isomerization may in some way modulate Abc2 activity and therefore P22 infection. In this respect, it is interesting to note that proline isomerization has been found to be critical for infection of *E. coli* by phage fd (47). Further work will be required to fully understand the influence of P22 infection on homologous recombination in bacterial cells which may also have implications of our understanding of RecA loading by the native RecBCD complex.

The RecBCD complex and related systems such as AddAB (also known as RexAB) have been considered as attractive targets for anti-bacterial drug development (48, 49). This reflects the fact that loss of RecBCD/AddAB function reduces infectivity (50–52), potentiates the effects of DNA-damaging agents (10, 53–56), and reduces the potential for adaptive mutagenesis leading to anti-bacterial resistance (54, 57, 58). These effects are all caused by a failure to efficiently respond to DNA breakage caused either by host immune responses or treatment with drugs such as fluoroquinolones. The diverse mechanisms employed by phage to manipulate bacterial DNA break repair may provide inspiration for the ongoing design of small molecule inhibitors for RecBCD/AddAB (48, 56).

## Acknowledgements

Work in the MSD laboratory was funded by the Wellcome Trust (100401/Z/12/Z) and the BBSRC (BB/S007261/1). Work in the DBW laboratory was funded by Cancer Research UK (C6913/A2160), the Wellcome Trust (209327/Z/17/Z) and the MRC (MR/N009258/1). We thank Martin Webb (The Crick Institute) for the gift of the SSB biosensor and Emma Galletti di Cadilhac (Bristol) for technical assistance.

## Supplementary Information

### Supplementary Figure Legends

**Supplementary Figure 1.**
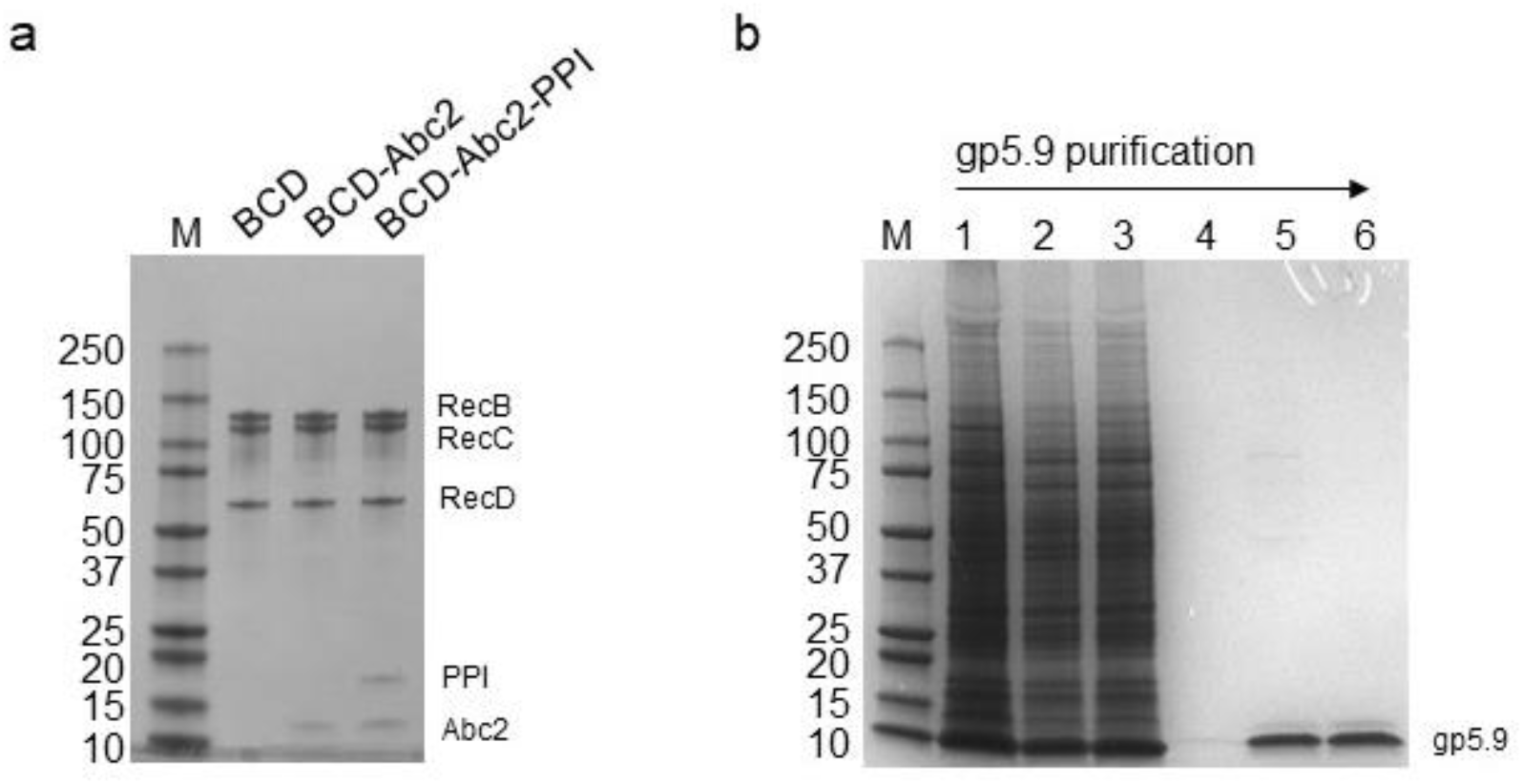
Purified proteins and protein complexes used in this study. (A) Purified RecBCD, RecBCD-Abc2^P68A^ and RecBCD-PPI-Abc2 complexes (1.5 μg each) were run on SDS-PAGE gels. The subunits are indicated. (B) Purification of gp5.9 from an insect cell extract. Lane 1, whole cell extract; lane 2, soluble cell extract; lane 3, Ni^2+^ flowthrough; lane 4, Ni^2+^ elution; lane 5, monoQ elution, lane 6; final prep after SEC and concentration.

**Supplementary Figure 2.**
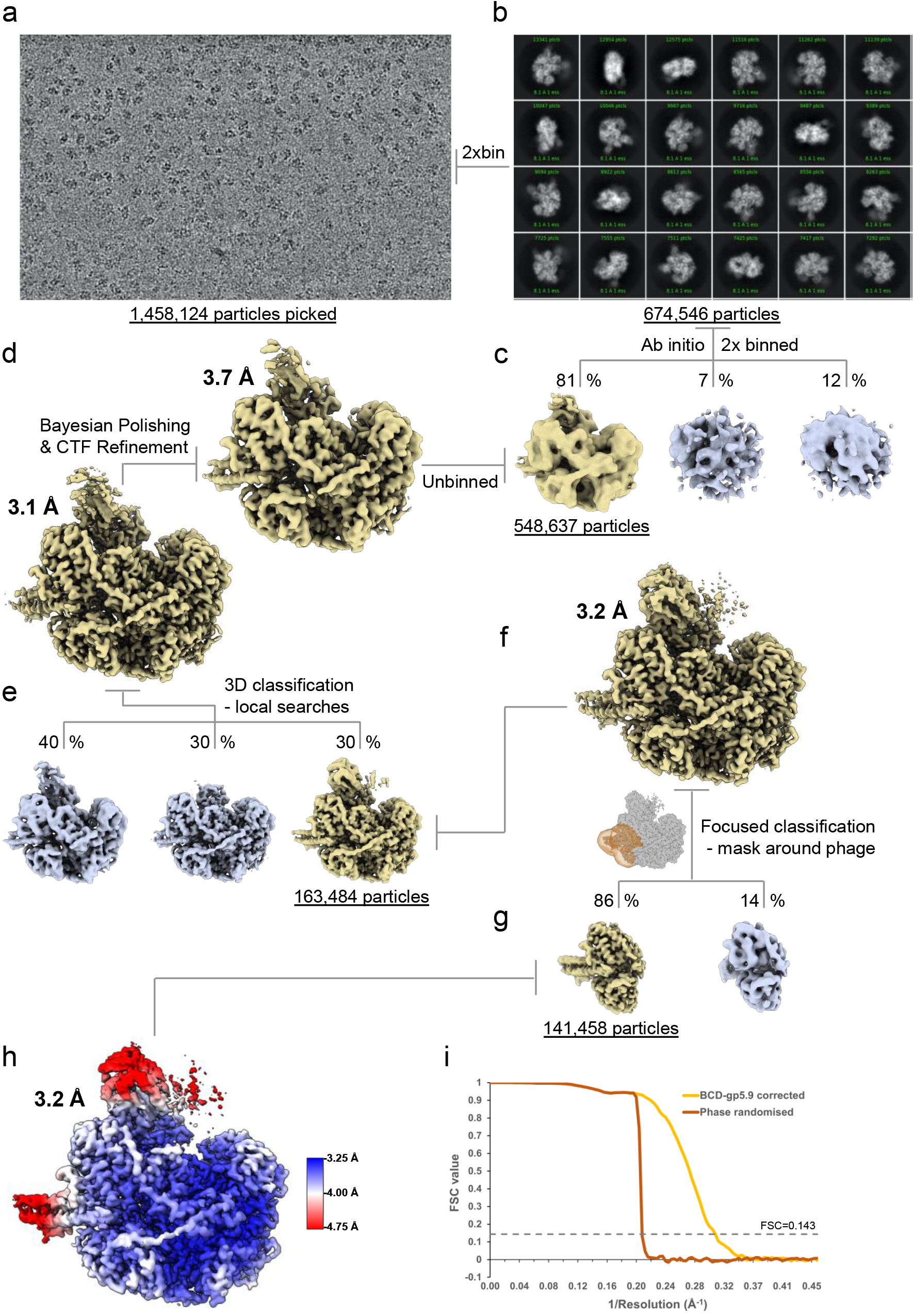
CryoEM processing scheme for the RecBCD-gp5.9 dataset. (A) Representative micrograph from the dataset. (B) The most populated 2D class averages generated in cryoSPARC from the cleaned 2xbinned particle set. (C) The results of cryoSPARC ab initio 3D classification. (D) The RELION-3 refined cryoEM maps before and after Bayesian polishing and CTF refinement. (E) The output classes from 3D classification with local angular searches. (F) 3D refinement of the selected particles and generation of a mask around the gp5.9 binding site used for (G) focused classification to remove particles absent of density for gp5.9. (H) The final refined RecBCD-gp5.9 map coloured by local resolution as estimated in RELION. (I) FSC curves for the corrected and phase randomised maps.

**Supplementary Figure 3.**
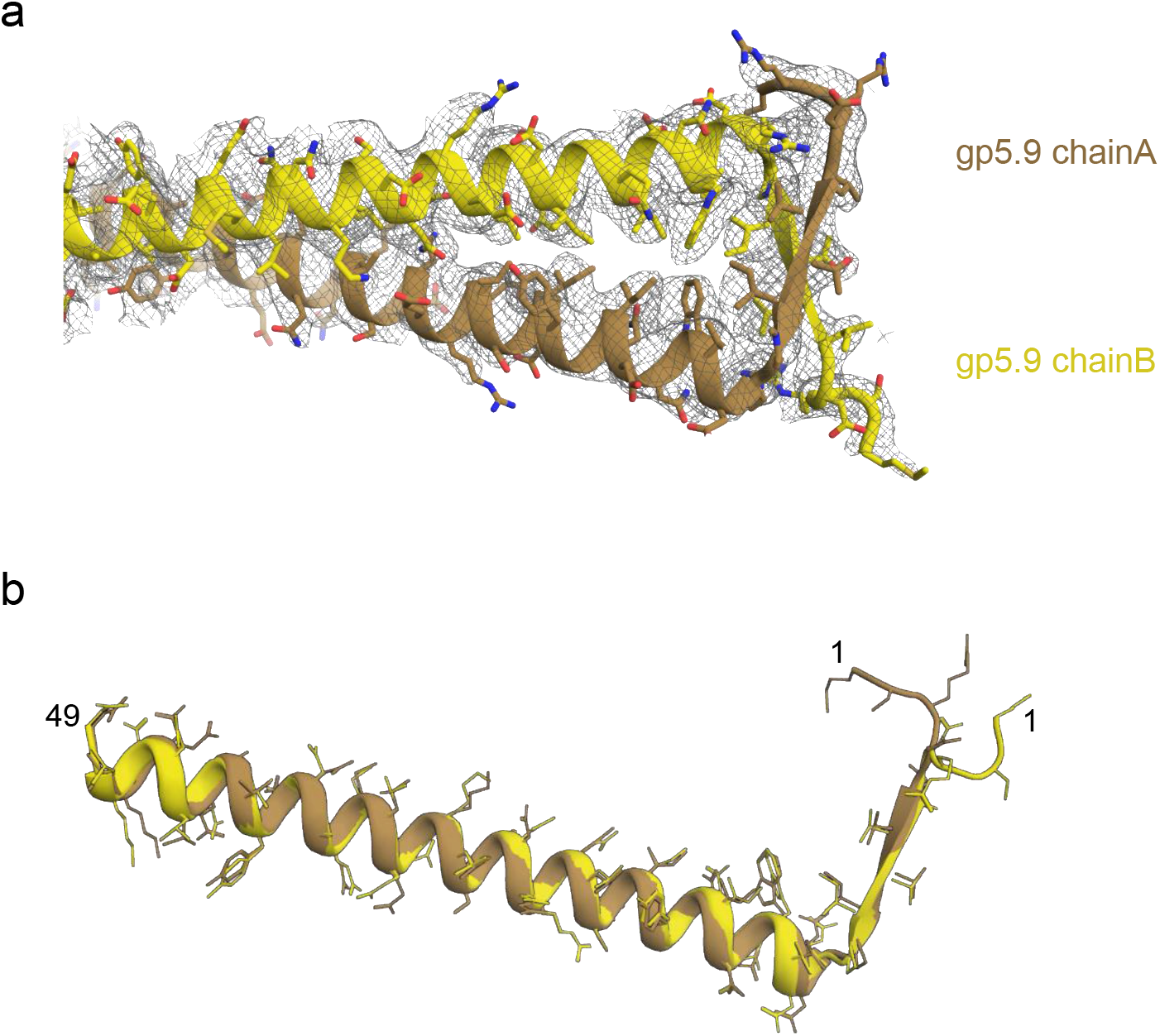
Modelling the structure of the gp5.9 phage protein. (A) The model of the gp5.9 dimer with the surrounding cryoEM density represented by grey mesh. (B) Superposition of the two chains of the gp5.9 model shows they share a high degree of structural similarity.

**Supplementary Figure 4.**
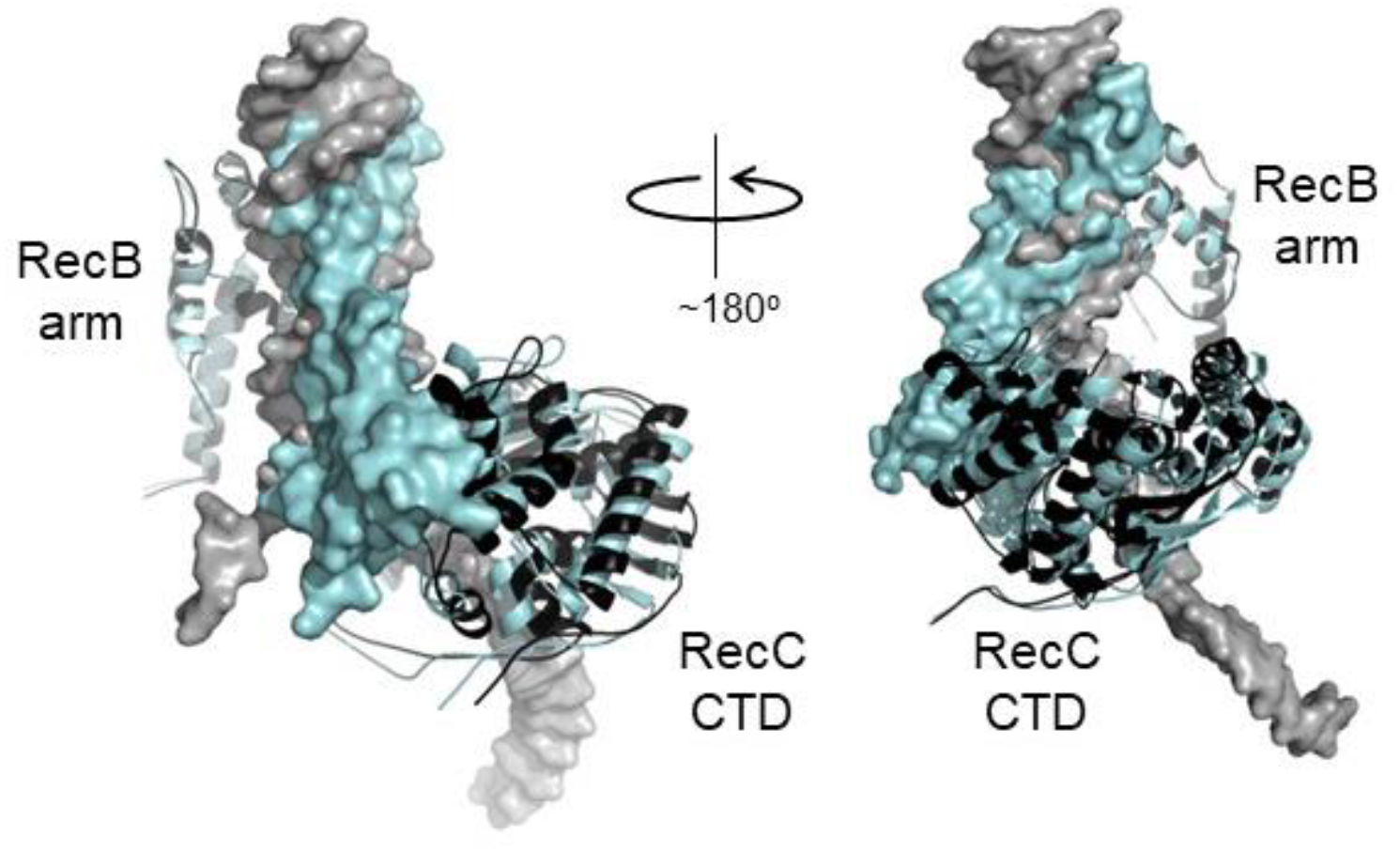
Small rigid body domain movements in RecBCD facilitate binding to either DNA or gp5.9. Both DNA and gp5.9 bind predominantly to RecBCD at the RecB arm domain and the RecC CTD. Small changes in the relative positioning of these two domains allow RecBCD to bind either DNA or gp5.9 at the same site despite their slightly different sizes and orientation of binding. The RecBCD-DNA complex and the RecBCD-gp5.9 complex are shown overlaid and in black and cyan respectively. In both cases the ligand (i.e. either DNA or gp5.9) is shown as a surface representation whereas the RecB arm and RecC CTD are shown as ribbons (positions indicated). The structures have been aligned using the RecB arm domains which are therefore almost perfectly superimposed upon one another. Note the different positioning of the RecC CTD which accommodates differences in the binding of either DNA or the DNA mimic.

**Supplementary Figure 5.**
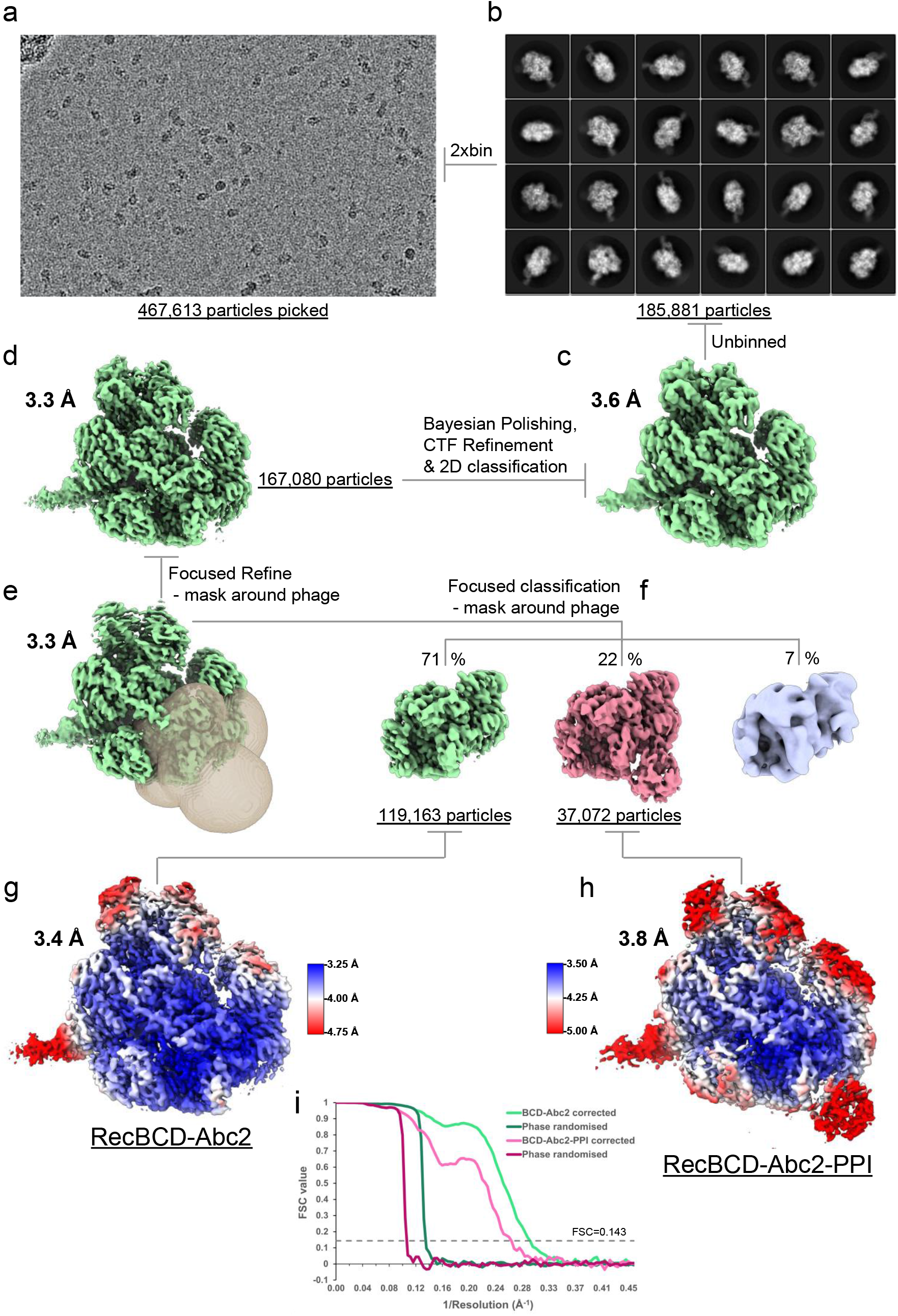
CryoEM processing scheme for the RecB^D^CD-Abc2-PPI-DNA dataset. (A) Representative micrograph from the dataset. (B) The most populated 2D class averages generated in cryoSPARC from the cleaned 2xbinned particle set. The RELION-3 refined cryoEM maps (C) before and (D) after Bayesian polishing and CTF refinement. (E) Resulting map from a focused 3D refinement using a mask around the Abc2 binding site with the mask displayed as a transparent surface. (F) Focused classification isolated a high resolution RecBCD-Abc2 class and a RecBCD-Abc2-PPI class. The final refined (G) RecB^D^CD-Abc2-DNA and (H) RecB^D^CD-Abc2-PPI-DNA map coloured by local resolution as estimated in RELION. (I) FSC curves for the corrected and phase randomised maps.

**Supplementary Figure 6.**
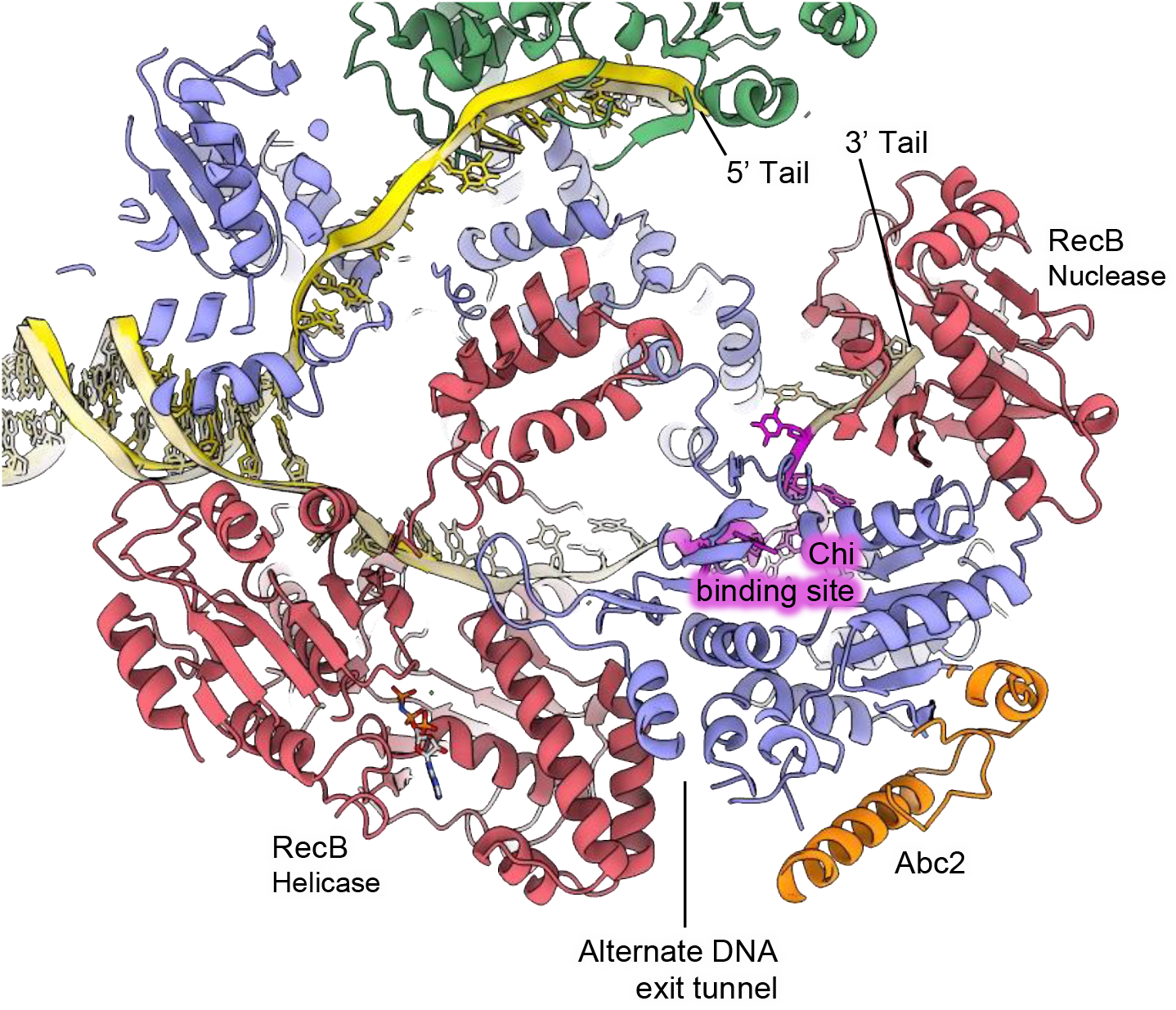
Overview of the location of Abc2 binding in the context of the RecBCD complex. Slabbed view of the RecBCD-Abc2-DNA structure with the Chi DNA substrate from the RecBCD Chi recognition CryoEM structure (PDB: 6SJB; (24)) overlaid in gold with the Chi residues highlighted in magenta.

**Supplementary Figure 7.**
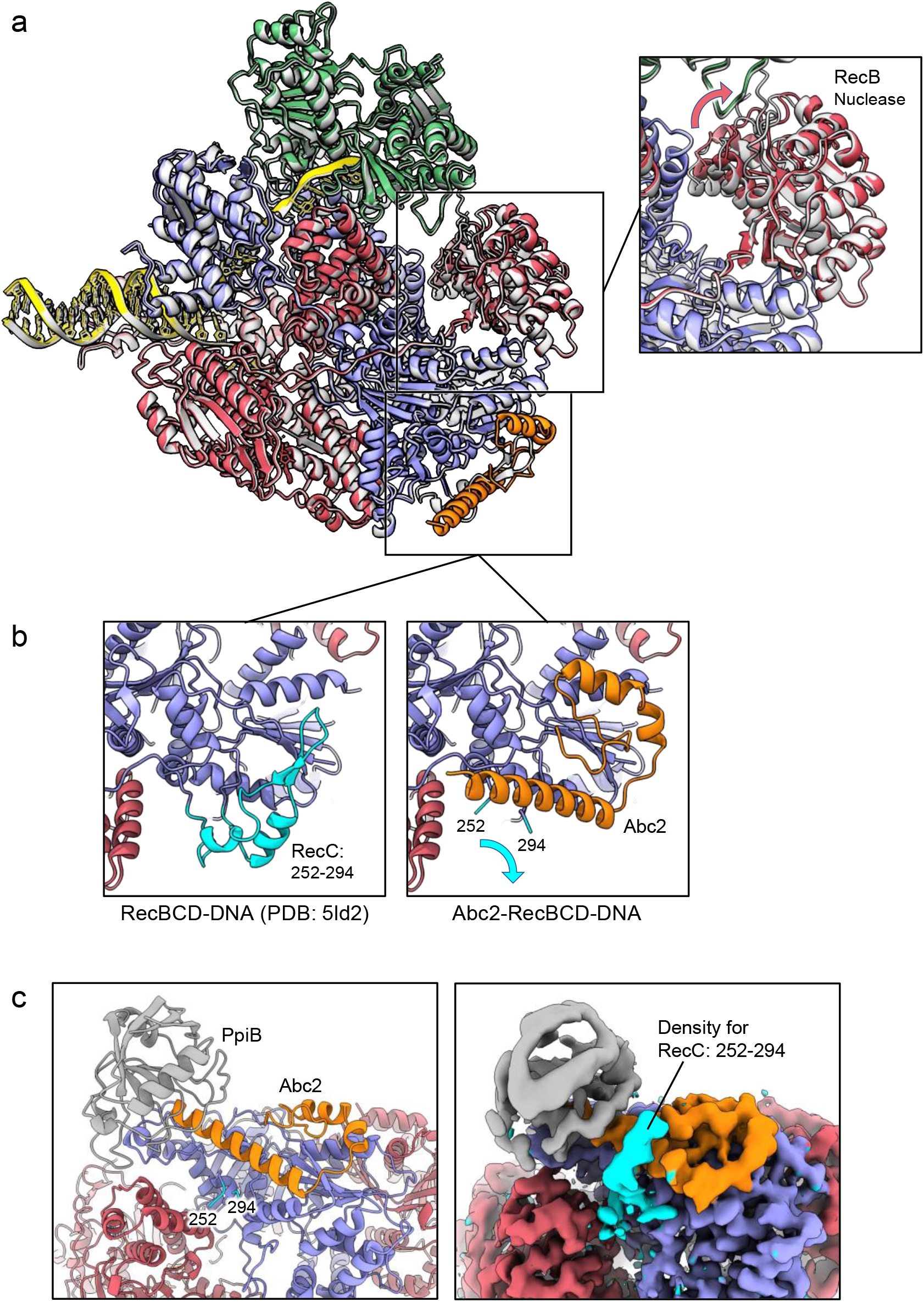
Abc2 binding induces minimal conformational changes in the RecBCD complex. (A) Superposition of the RecBCD-Abc2-DNA structure (coloured as in other figures) with the corresponding RecBCD-DNA structure containing the same DNA substrate (PDB: 5LD2) in grey. The boxed panel shows a close-up of the superposition, revealing a slight rotation within the RecB nuclease domain. (B) Side-by-side related close-up views of the Abc2 binding site in the structures without (left) and with (right) Abc2. A surface helical bundle involving RecC^252-294^, highlighted in cyan, peels back to allow Abc2 binding. (C) Rotated view of the Abc2 binding site in the RecBCD-Abc2-PPI-DNA model (left) with the associated cryoEM map displayed unsharpened (right) to indicate a potential path for the displaced RecC^252-294^ domain, highlighted in cyan.

**Supplementary Figure 8.**
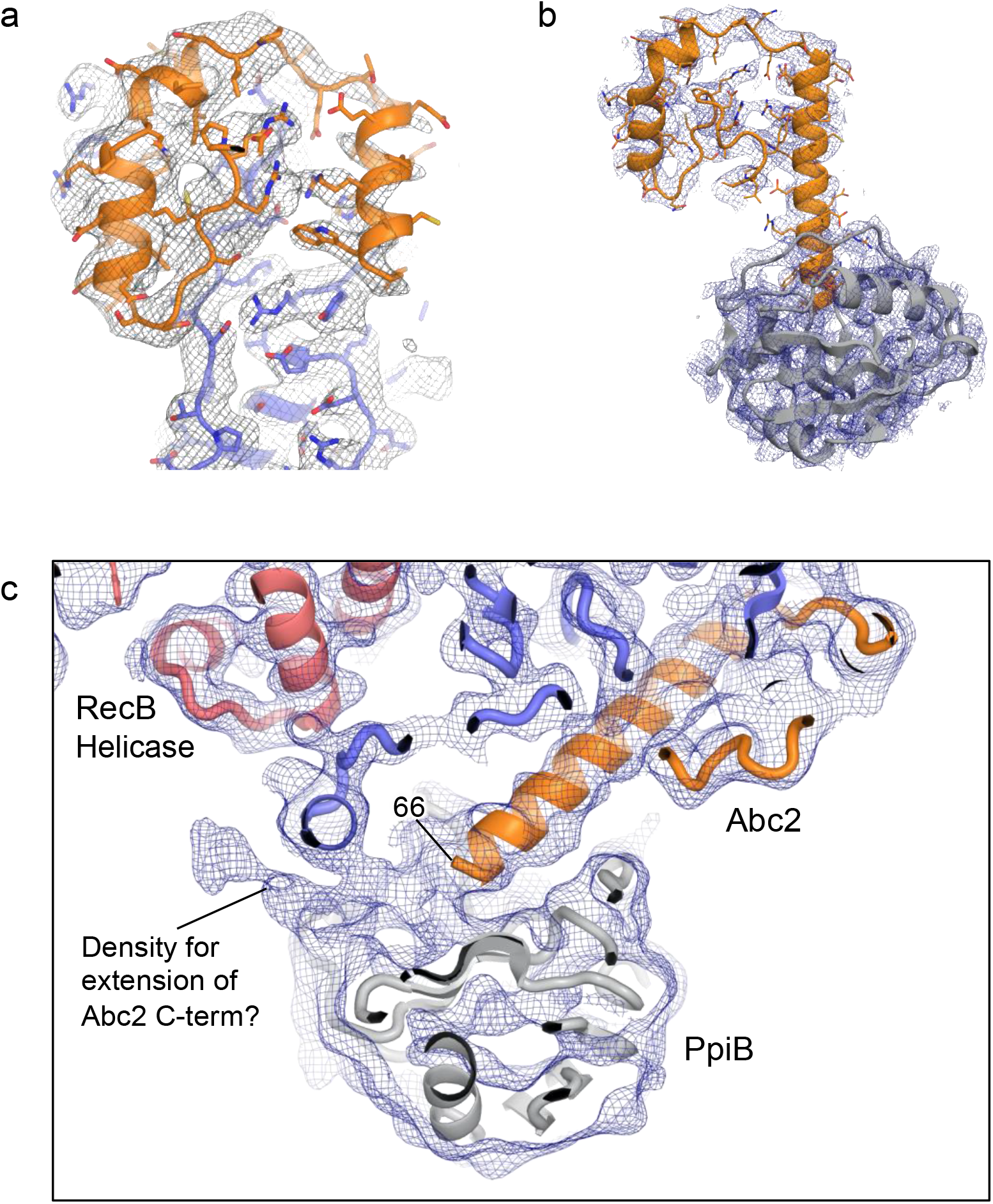
Modelling of Abc2 and PpiB in the CryoEM maps. (A) Density and model for the RecBCD-Abc2-DNA structure showing the Abc2 binding site on RecC. (B) Density and model for Abc2 and docked PpiB taken from the RecBCD-Abc2-PPI-DNA structure. (C) View of the blurred RecBCD-Abc2-PPI-DNA map showing evidence for the continuation of the Abc2 C-terminus through the crevice between PpiB and RecC.

**Supplementary Figure 9.**
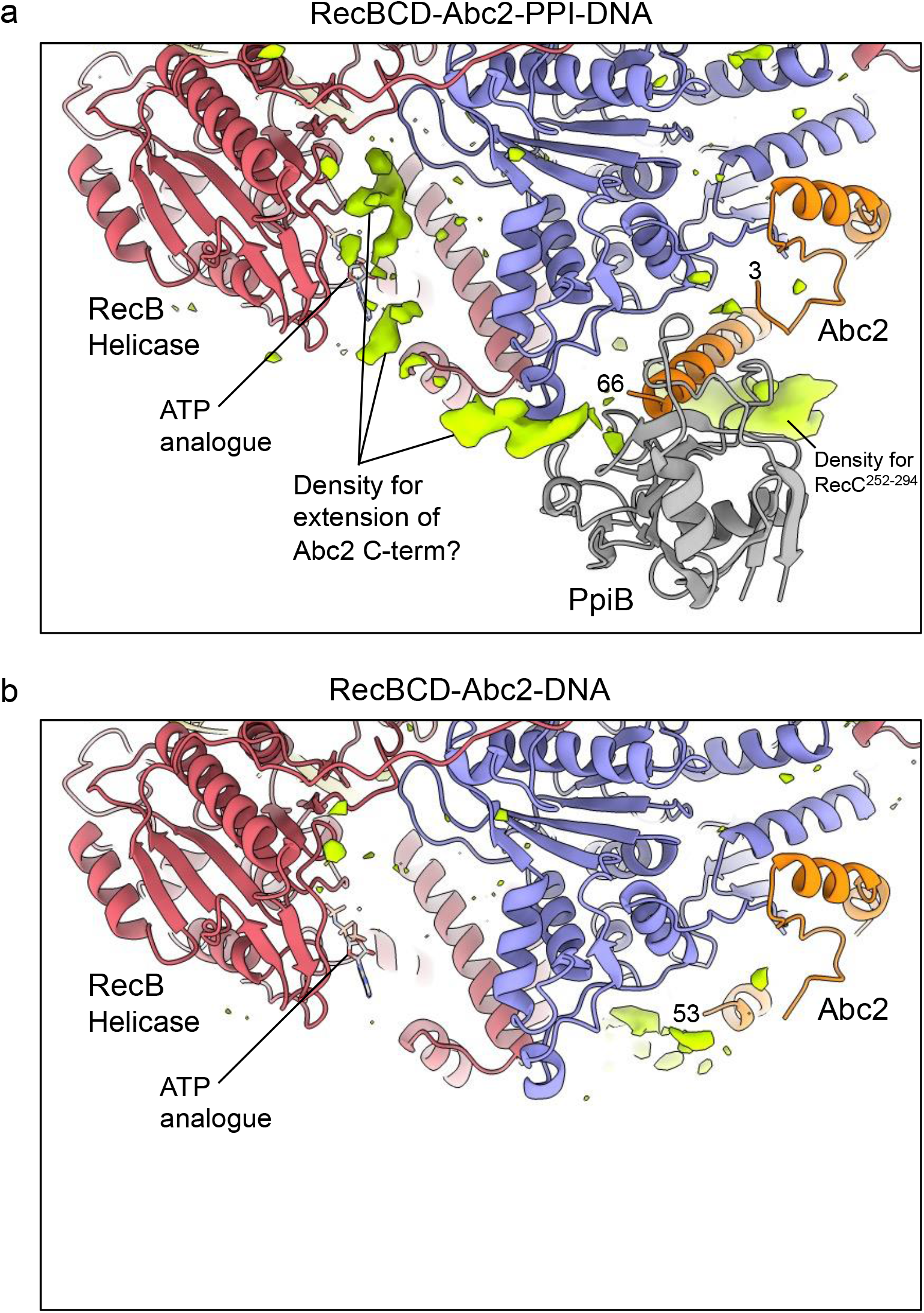
Additional unmodelled density suggests a path for the Abc2 C-terminus breaching the RecB helicase domains. Similar views and contouring the of blurred (A) RecBCD-Abc2-PPI-DNA and (B) RecBCD-Abc2-DNA CryoEM maps with additional, unmodelled difference density shown as a lime green surface. This suggests that in the PpiB bound complex the C-terminus of Abc2 becomes partially ordered around the RecB helicase domains, in the vicinity of the bound ATP analogue ADPNP.

**Supplementary Figure 10.**
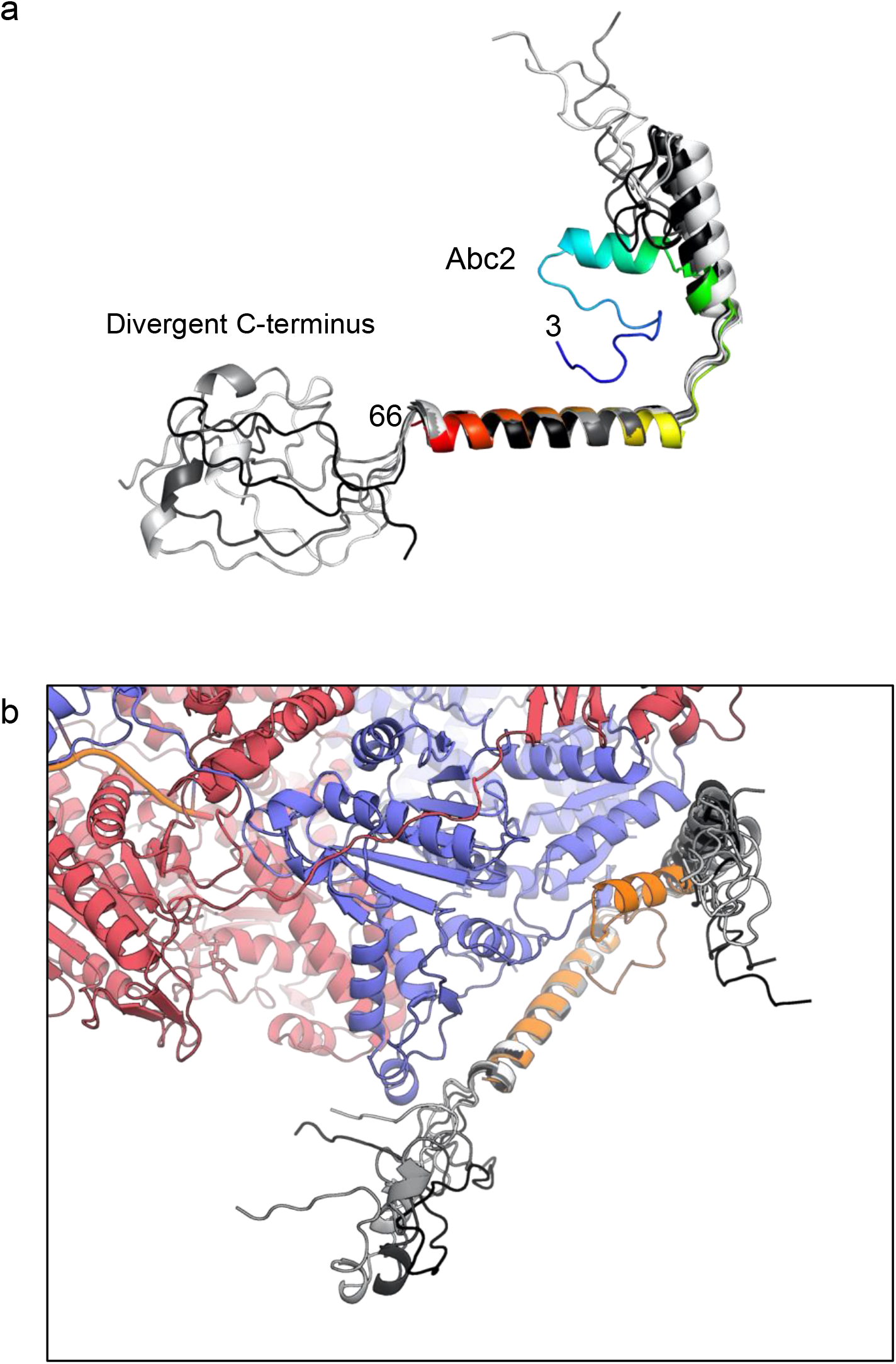
AlphaFold modelling of the full Abc2 protein shows a divergent C-terminus. The five top AlphaFold models for the Abc2 protein (coloured on a greyscale spectrum) are aligned to the solved Abc2 model (residues 3-66, coloured blue-to-red from the N- to the C-terminus). The predictions diverge significantly in trying to assign structure to the C-terminal residues which cannot be modelled in the CryoEM structure. (B) Alignment of the five AlphaFold models to Abc2 in the context of the RecBCD-Abc2-DNA structure. None of the predictions orientate the C-terminus is a way that would satisfy the additional density seen within the RecB helicase.

**Supplementary Figure 11.**
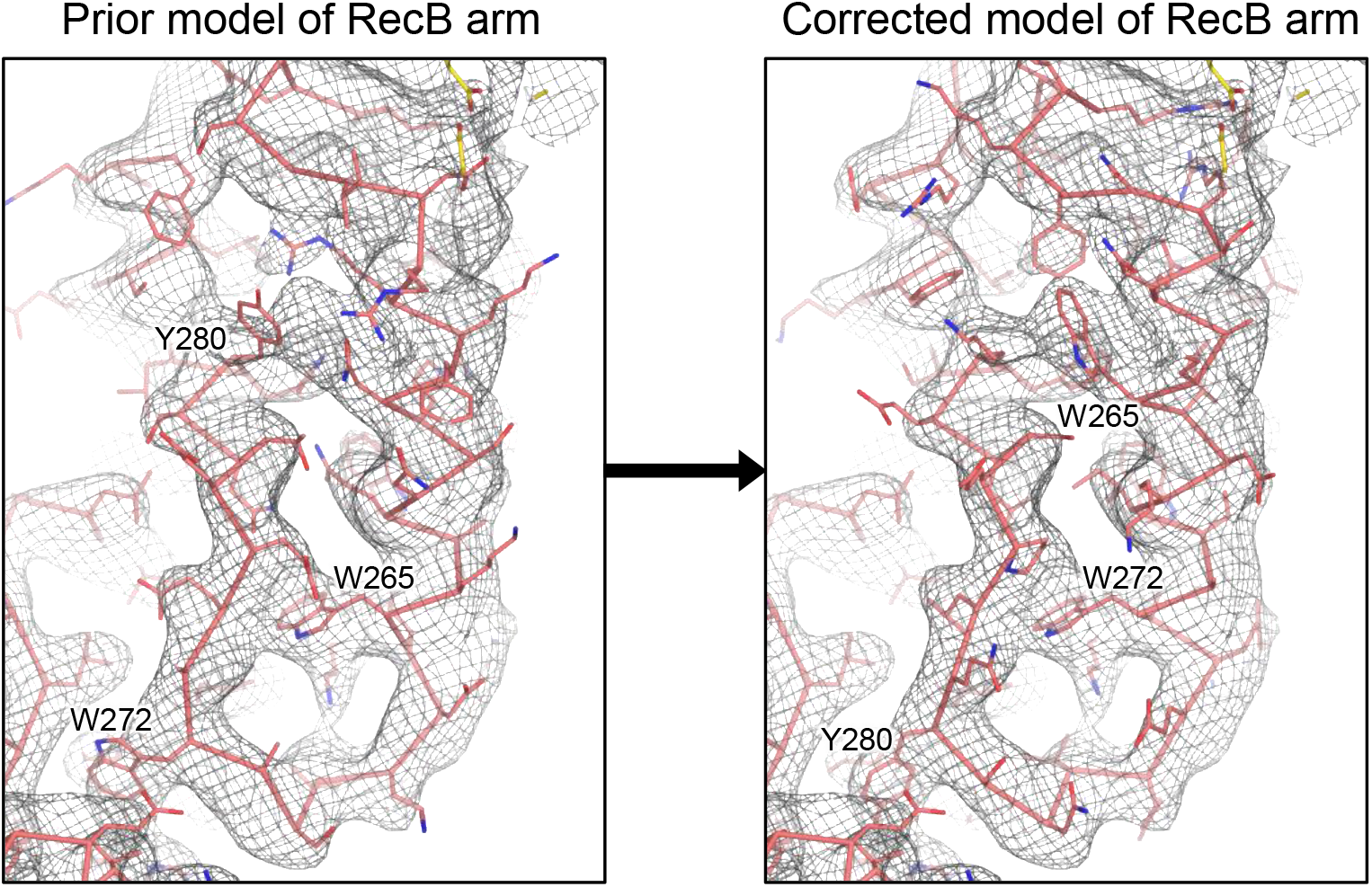
Rebuilding of the RecB arm domain model based on the high resolution RecBCD-gp5.9 CryoEM map. Side-by-side view of the density for the RecB arm in the RecBCD-gp5.9 CryoEM map, with the original RecB model (left) shown alongside the corrected RecB model (right). The large hydrophobic residues key to establishing the register of the model are labelled, highlighting the 7-8 residue sequence shift induced by the rebuilding.

